# A SEVA-based, CRISPR-Cas3-assisted genome engineering approach for *Pseudomonas* with efficient vector curing

**DOI:** 10.1101/2023.06.29.547033

**Authors:** Eveline-Marie Lammens, Daniel Christophe Volke, Kaat Schroven, Marleen Voet, Alison Kerremans, Rob Lavigne, Hanne Hendrix

## Abstract

The development of CRISPR-Cas-based engineering technologies has revolutionized the microbial biotechnology field. Over the years, the Class II Type II CRISPR-Cas9 system has become the gold standard for genome editing in many bacterial hosts. However, the Cas9 system does not allow efficient genomic integration in *Pseudomonas putida*, an emerging Synthetic Biology host, without the assistance of lambda-Red recombineering. In this work, we utilize the alternative Class I Type I-C CRISPR-Cas3 system from *Pseudomonas aeruginosa* as a highly-efficient genome editing tool for *P. putida* and *P. aeruginosa*. This system consists of two vectors, one encoding the Cas genes, CRISPR array and targeting spacer, and a second SEVA-vector, containing the homologous repair template. Both vectors are Golden Gate compatible for rapid cloning and are available with multiple antibiotic markers, for application in various Gram-negative hosts and different designs. By employing this Cas3 system, we successfully integrated an 820-bp cassette in the genome of *P. putida* and performed several genomic deletions in *P. aeruginosa* within four days, with an efficiency of >83% for both hosts. Moreover, by introducing a universal self-targeting spacer, the Cas3 system rapidly cures all helper vectors, including itself, from the host strain in a matter of days. As such, this system constitutes a valuable engineering tool for *Pseudomonas*, to complement the existing range of Cas9-based editing methods and facilitates genomic engineering efforts of this important genus.

**Importance:** The CRISPR-Cas3 editing system as presented here facilitates the creation of genomic alterations in *P. putida* and *P. aeruginosa* in a straightforward manner. By providing the Cas3 system as a vector set with Golden Gate compatibility and different antibiotic markers, as well as by employing the established SEVA vector set to provide the homology repair template, this system is flexible and can readily be ported to a multitude of Gram-negative hosts. Besides genome editing, the Cas3 system can also be used as an effective and universal tool for vector curing. This is achieved by introducing a spacer that targets the *oriT*, present on the majority of established (SEVA) vectors. Based on this, the Cas3 system efficiently removes up to three vectors in only a few days. As such, this curing approach may also benefit other genomic engineering methods or remove naturally-occurring plasmids from bacteria.

## Introduction

The *Pseudomonas* genus comprises a variety of aerobic, Gram-negative bacteria that are ubiquitous in nature, playing diverse biological roles that range from plant growth promotion to bioremediation and pathogenicity (1). The genus contains over 140 species (www.catalogueoflife.org) and is one of the most ecologically and medically important groups of bacteria. This includes the well-known antibiotic resistant pathogen *Pseudomonas aeruginosa*, which is responsible for infections in immunocompromised patients (2), and the biochemical versatile species *Pseudomonas putida* involved in industrial processing (3). Therefore, robust engineering tools for *Pseudomonas* cannot only support fundamental discoveries, but also modify pathogenicity, improve production yield, and enable development of microbial cell factories (4, 5).

Diverse engineering systems are available to modify the genomes of *Pseudomonas* species. While the transposon-based systems insert DNA sequences in a random (e.g. Tn5 transposon) or site-specific manner (e.g. Tn7 transposon) and cannot delete genes (only disrupt them) (6, 7), the homologous recombination methods with integrative plasmids, such as the two-step allelic exchange and I-SceI-mediated recombination, require two rounds of selection using chromosomal markers with often low recombination frequency to achieve scar-less genome editing (8, 9). More efficient recombineering methods using heterologous recombinases which catalyze recombination between similar sequences (e.g. λ Red and RecET recombinase systems) or between specific recognition sites (e.g. Cre/lox and Flp/FRT systems) also involve an additional step for integrated selection marker removal, extending the engineering time and often leaving a scar behind in the genome (10–13).

In recent years, the CRISPR-Cas (<u>c</u>lustered regularly *i*nterspaced <u>s</u>hort <u>p</u>alindromic repeats and <u>C</u>RISPR-<u>as</u>sociated proteins) systems have been proven to efficiently engineer genomes in virtually all species (14–19). The systems comprise an RNA guide called CRISPR RNA (crRNA) with sequence complementary to the target DNA (spacer), guiding a nuclease to make a site-specific double-strand break. Since many bacteria lack non-homologous end-joining to repair this break, a DNA repair template has to be provided to restore the defect by homologous recombination (20). Alternatively, the CRISPR-Cas system can be used as a counter-selection tool after recombineering or homologous recombination (21). Similar to other organisms, the most well-known engineering systems for *Pseudomonas* are based on single subunit Class 2 CRISPR systems. These include the Type II CRISPR-Cas9 system of *Streptococcus pyogenes* (*Sp*Cas9) (22–28) and the Type V CRISPR-Cas12a from *Francisella novicida* (*Fn*Cas12a) (23, 29). However, these systems do not allow genomic integration of sequences based on a repair template in *P. putida* (23). The Class 1 Type I system CRISPR-Cas3, on the other hand, consists of a multi-subunit complex and has the advantage to be the most prevalent in nature, enabling engineering with endogenous systems, and to degrade DNA processively, allowing larger deletions (30, 31). Recently, Csörgő et al. (2020) exploited the Type I-C CRISPR-Cas3 system from *P. aeruginosa* (*Pae*Cas3c) for heterologous genome engineering in various microbial species, obtaining genome-scale deletions with random and programmed size and recombination efficiencies surpassing those of the *Sp*Cas9-based system. Moreover, CRISPR-Cas3 has been introduced as the base editing tool CoMuTER, for targeted *in vivo* mutagenesis in yeast (32).

One major hurdle of CRISPR-Cas-assisted methods as well as other commonly used engineering techniques is the use of auxiliary plasmids, which need to be removed from the bacterial cells after engineering. Well-known curing systems rely on counter-selectable markers, repeated passaging of the cells, the use of tractable vectors, DNA intercalating agents or conditional origins-of-replication (33–37). Nevertheless, these methods are often time-consuming, laborious, not effective in some bacteria, can introduce off-target genomic mutations or require specific vectors and conditions for their functionality (37–41). To avoid these issues, CRISPR-Cas-based plasmid curing systems showed to be promising. Indeed, a recently developed CRISPR-Cas9-assisted curing system (pFREE) showed efficiencies between 40 and 100% for the major classes of vectors used in molecular biology, including SEVA vectors, by targeting conserved sequences within origins-of-replication in multiple bacterial backgrounds (42, 43).

In this study, an efficient scar-less genome editing and plasmid curing method based on CRISPR-Cas3 was developed for *Pseudomonas*. The system consists of the all-in-one pCas3cRh targeting plasmid designed by Csörgő et al. (2020) combined with the Standard European Vector Architecture (SEVA) vectors for homologous directed repair and curing, resulting in a straightforward, efficient and universal system for genomic deletion and integration. The applicability of the method is demonstrated in *P. putida* KT2440 and SEM11 and *P. aeruginosa* PAO1. Moreover, the system has been expanded by making it Golden Gate compatible, adding several antibiotic markers and including a fluorescent marker to facilitate the screening procedure.

## Results and Discussion

### An overview of the CRISPR-Cas3-based engineering approach for *Pseudomonas*

A CRISPR-Cas3-based engineering method was developed, which enables the creation of genomic deletions, insertions or substitutions in the *Pseudomonas* genome in an efficient and flexible manner. In general, the Cas genes (*cas3, cas5, cas7* and *cas8*) and crRNA with spacer sequence are all located on the pCas3cRh vector under the control of the RhaRS/*P*_*RhaBAD*_ inducible system. Guided by the crRNA, the Cas3 enzyme creates a targeted cut in the genomic DNA upon induction with rhamnose. After cleavage, the damaged genome will be restored by homology-directed repair (HDR) to create the desired genomic modification. To perform the HDR, a homology repair template is provided on vector pSEVA231 (KmR, for *P. putida*) or pSEVA131 (CbR, for *P. aeruginosa*). The design of the repair template determines the prospective modification of the genome, namely a deletion, insertion of substitution. It is important to note that any canonical SEVA vector can serve as a carrier for the repair template, which allows the user to select a backbone with his or her preferred antibiotic marker and origin of replication for the application in mind and allows compatibility with any Gram-negative host (44).

Finally, after verification of the correct genomic modification with PCR and sequencing, the pCas3cRh and pSEVAX3-HDR vectors are cured from the host by introduction of pSEVA52-oriT. This vector expresses a spacer sequence targeting the *oriT* (origin-of-transfer) site, which is located on all SEVA plasmids (including itself) as well as many other established vectors and will enable the swift restriction and removal of the helper vectors (Figure 1).

**Figure 1:**
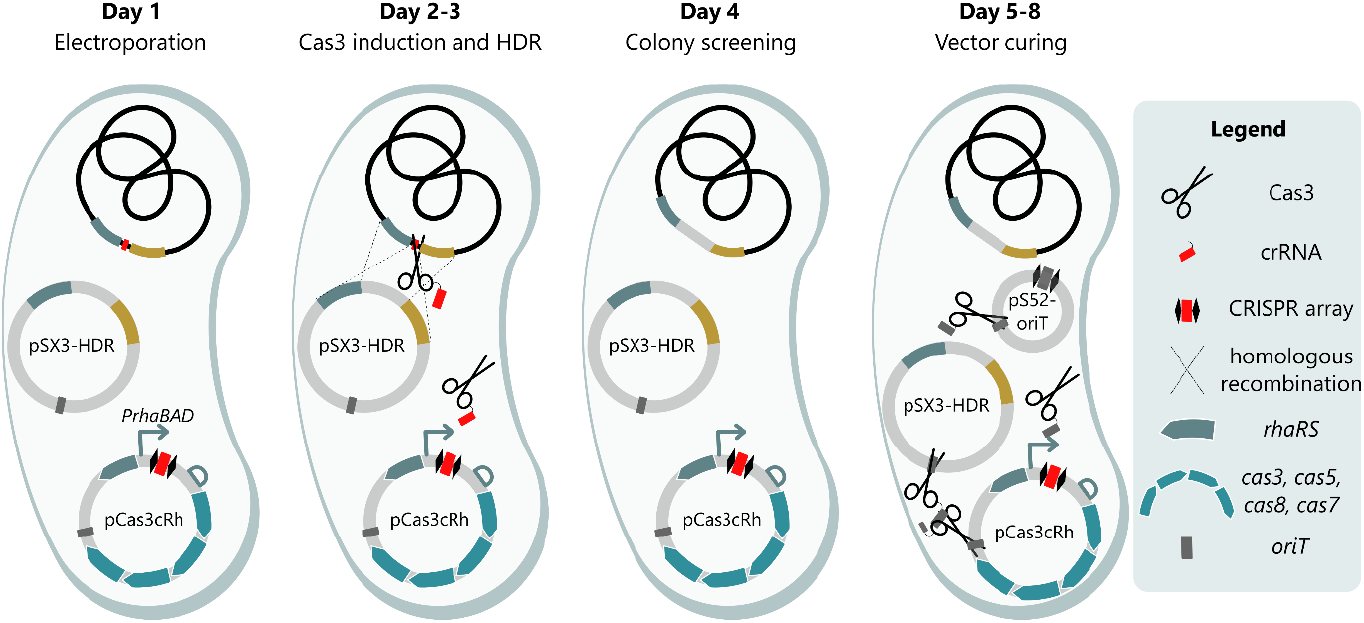
General overview of the CRISPR-Cas3-based engineering method in *Pseudomonas*, as illustrated for *P. putida*. Day 1: The pCas3cRh vector with spacer sequence (red) for the target site and pSEVA231 with the repair template for homology-directed repair (pSX3-HDR) are introduced in *Pseudomonas* by means of electroporation. Day 2-3: On day 2, multiple colonies are inoculated together in growth medium supplemented with rhamnose to induce expression of the Cas3 system. The Cas3 enzyme cleaves the genomic DNA at the target location, i.e. the protospacer (marked in red), after which the dsDNA break will be repaired by homologous recombination using the repair template. After overnight induction, a dilution streak on LB agar is performed on day 3. Day 4: Multiple single colonies of the dilution streak are analyzed by PCR and Sanger sequencing, to verify the presence of the desired genomic modification. Correct mutants are grown overnight to start the vector curing process. Day 5-8: On day 5, the overnight cultures are transformed with pSEVA52-oriT (pS52-oriT), which carries a spacer sequence targeting the origin-of-transfer (*oriT*) broadly used on plasmids. Similar to day 2-3, expression of the Cas3 system is induced on day 6, which will cleave all vectors and lead to efficient curing of the host strain. After a dilution streak on day 7 and overnight incubation, correct vector curing is verified on day 8 by streaking individual colonies on all antibiotics separately that were used to select the vectors. A similar method is applied for *P. aeruginosa*. However, no rhamnose is required to induce the system, and pSEVA131 is used for homology directed repair.

### The CRISPR-Cas3-based engineering system enables efficient genomic engineering of *P. putida*

In the following section, the engineering method will be described and illustrated in detail by means of an integration example in *P. putida* KT2440 and *P. putida* SEM11. More specifically, an expression construct consisting of *P*_*14c*_*-BCD22-phi15lys(G3RQ)* is integrated in locus PP_5388 in both hosts, resulting in low, constitutive production of phi15 lysozyme (G3RQ) (Figure 2) (45). First, a PAM (protospacer adjacent motive) site is selected in proximity of the target, which will be the recognition site of the Cas3 enzyme. In general, the Cas3 system employs a 5’ AAG PAM with an upstream protospacer, however, in this work a TTC PAM is used in combination with a downstream protospacer consistent with the work of Csörgő *et al*. (2020). The PAM sequence is preferably located within the sequence that is to be deleted or substituted, or, in case of an integration, within 15 bp of the integration site. If no suitable PAM site is available in these regions, a site within the neighboring sequences of the genomic modification can be used as well, but the PAM site (or protospacer sequence) should be removed from the homology arms in later steps. The selected PAM site determines the spacer sequence, which is located directly downstream of the TTC trinucleotide, has a length of 34 bp and should not have significant homology to secondary sequences in the genome. The selected spacer sequence can be efficiently integrated in pCas3cRh by Golden Gate cloning with Type IIs restriction enzyme BsaI, as explained in the Method section. For the example for integration in PP_5388 in *P. putida*, a PAM site was selected 1 bp upstream of the intended integration site and the downstream spacer 5’-AGATCATGGTAACCCCGGCCGCTGGAGCCATTTC-3’ was successfully cloned into pCas3cRh to yield pCas3cRh-PP_5388 (Figure 2c) (Tables S1 and S2).

**Figure 2:**
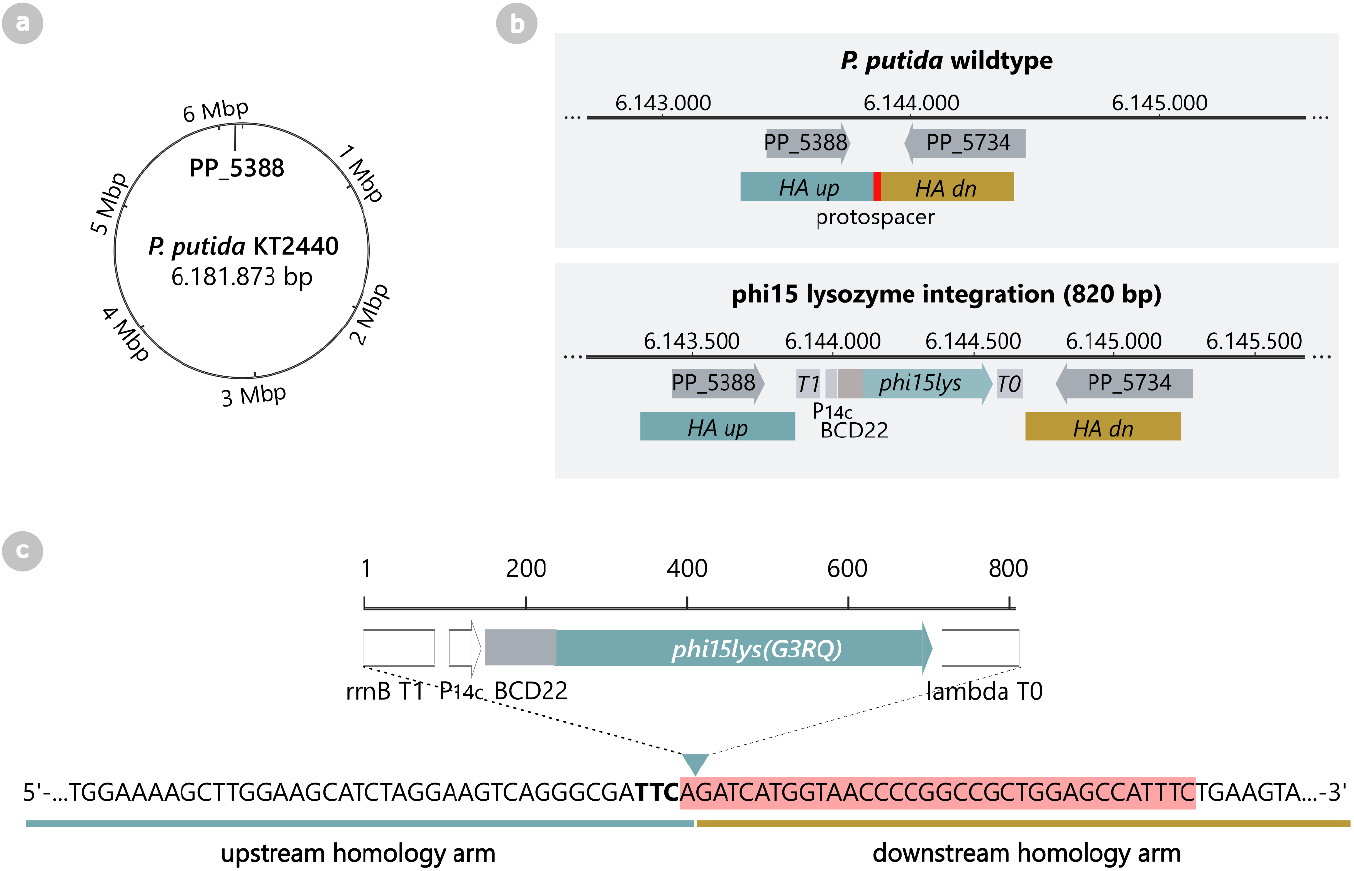
Genomic integration of expression cassette *P*_*14c*_*-BCD22-phi15lys(G3RQ)* in locus PP_5388 in *P. putida*. a) Genome of *P. putida* KT2440 with indication of locus PP_5388. b) Integration cassette *P*_*14c*_*-BCD22-phi15lys(G3RQ)* has a total length of 820 bp. This phi15 lysozyme mutant G3RQ was optimized to inhibit the activity of the T7-like RNA polymerase (RNAP) of phage phi15, to reduce basal expression of this RNAP in uninduced conditions, similarly to the established pET system (45). The PP5388 was previously identified as a locus that results in low expression levels of integrated sequences (46). As such, low levels of the phi15 lysozyme (G3RQ) will inhibit basal concentrations of the phi15 RNAP, while leaving sufficient active RNAP molecules upon induction of the pET-like system. c) The PAM site (indicated in bold) lies 1 bp upstream of the integration site (triangle) where the expression cassette will be integrated. The protospacer sequence (highlighted in red) is defined as the first 34 bp directly downstream of the PAM site.

After construction of the pCas3cRh-spacer vector, a second vector with the repair template needs to be assembled. Any canonical SEVA vector can be used for this purpose, however, for this work we selected pSEVA231 (*P. putida*) and pSEVA131 (*P. aeruginosa*) due to their medium-copy number origin and appropriate resistance marker for the respective isolates. For deletions, the repair template simply consists of two joined homology arms (≥500 bp each), identical to the sequences directly up- and downstream of the region to be deleted. For integration or substitution, on the other hand, the repair template consists of the desired insertion or substitution, flanked by the up- and downstream homology arms. It is important to note that selected recognition sites within the homology arms should be removed from the repair template, either by deletion, PAM mutation, or protospacer/PAM interruption. For the selected integration in PP_5388, homology arms of 550 bp each were amplified from the genome *of P. putida* and ligated to flank the integration cassette in pSEVA23-PP_5388, using Golden Gate cloning with Type IIs restriction enzyme BsaI (Figure 2b, Tables S1 and S2).

Following the vector construction, both the pCas3cRh-spacer and the template vector are simultaneously introduced into the *Pseudomonas* host by co-electroporation. If the efficiency of the co-electroporation is insufficient, the vectors can be introduced consecutively by first introducing the repair template followed by pCas3cRh-spacer. For the PP_5388 integration, both *P. putida* KT2440 and *P. putida* SEM11 were successfully co-transformed and no morphological differences of the colonies were observed in comparison to electroporation with empty control vectors. Furthermore, pCas3cRh-PP_5388 was also successfully introduced separately, indicating that little to no basal expression occurs from the Cas3 system in *P. putida* and that the RhaRS/*P*_*rhaBAD*_ expression system is tightly regulated. To confirm this, 24 co-transformants of *P. putida* KT2440 and *P. putida* SEM11 analyzed by PCR with primers binding on the genome outside the homology arms, showing that none of the co-transformants had the desired insertion before induction of the CRISPR-Cas3 system (Figure S1).

To induce the CRISPR-Cas3 system, several co-transformants were pooled and used to inoculated 20 mL LB medium with the required antibiotics and 0.1% rhamnose. The cultures were then incubated overnight at the appropriate temperature. The following day, a dilution streak of the induced overnight culture was performed on agar plates with the appropriate antibiotics and grown until visible colony formation the following day. For the PP_5388 integration example, again 24 colonies of *P. putida* KT2440 and *P. putida* SEM11 were subjected to PCR with primers binding outside the homologous arms on the genome. Interestingly, after induction with rhamnose, 83% of *P. putida* KT2440 colonies and 88% of *P. putida* SEM11 colonies showed an amplicon length correlating to correct integration of the *P*_*14c*_*-BCD22-phi15lys(G3RQ)* cassette (Figure 3a, Figure S2). In comparison for uninduced control samples, no integration was observed in any of the screened *P. putida* KT2440 or *P. putida* SEM11 colonies (Figure 3a, Figure S2). As such, the Cas3 system is able to efficiently perform genomic integrations in *P. putida* without the assistance of any recombineering genes as required for the Cas9 system (23, 47).

**Figure 3:**
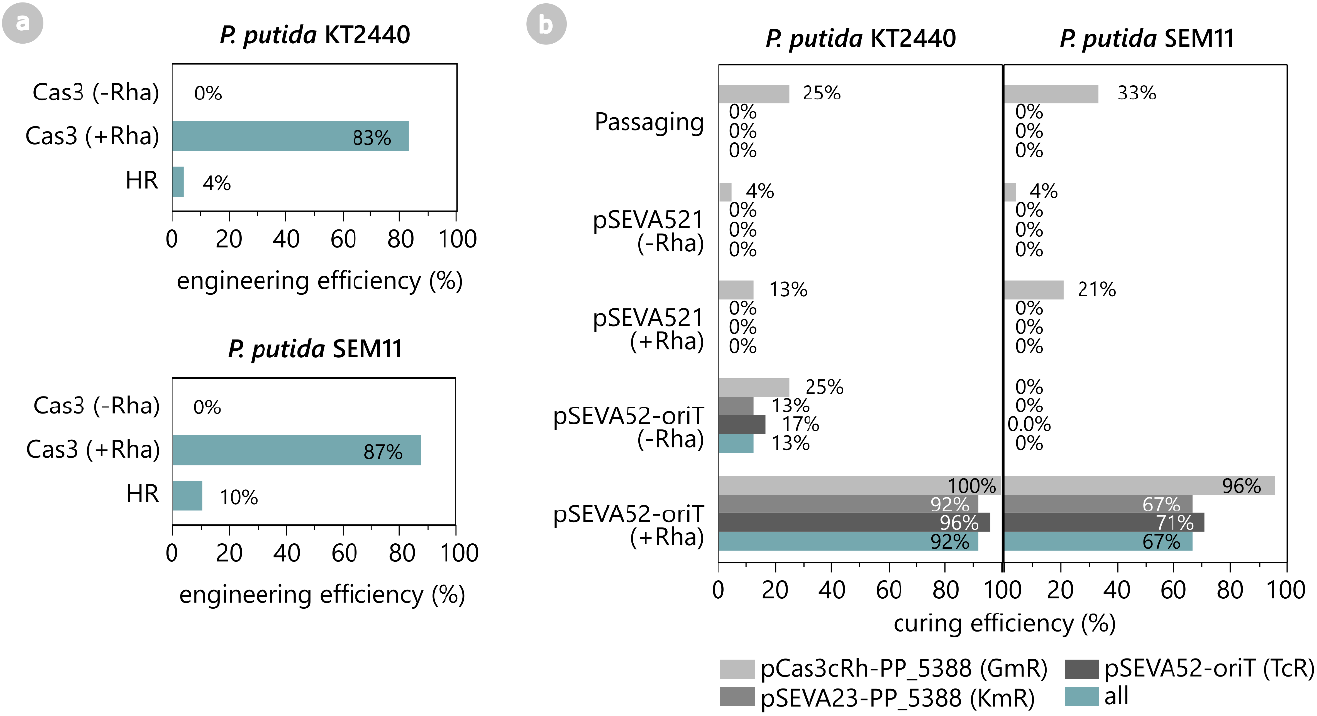
a) Engineering efficiencies of the integration of expression cassette *P*_*14c*_*-BCD22-phi15lys(G3RQ)* in locus PP_5388 of *P. putida* KT2440 and *P. putida* SEM11 using the CRISPR-Cas3-based method (Cas3), with or without induction with rhamnose (+/- Rha), or via traditional homologous recombination (HR). b) After engineering of *P. putida* with pCas3cRh-PP_5388 and pSEVA23-PP_5388, the strains are cured from the engineering vector by serial passaging or by CRISPR-Cas3-based curing using pSEVA52-oriT, with or without rhamnose induction (+/-Rha). As a negative control for the CRISPR-Cas3-based curing, an empty pSEVA521 vector was used instead of pSEVA52-oriT.

To put these engineering efficiencies into perspective, the same integration in *P. putida* was created using traditional homologous recombination (HR). More specifically, the two-vector system as described by Volke *et al*. (36) was employed, where the first vector, carrying the homology arms and desired modification, fully integrates in the genome in a first HR event. This event can be tracked by a green fluorescent reporter and antibiotic resistance marker on the integration vector. Next, a second vector supplies the I-SceI restriction enzyme, which will recognize and cut a unique restriction site within the integrated vector and force the second HR event, resulting in the desired genomic modification with loss of the fluorescent reporter and antibiotic marker. In this work, we successfully constructed integration vector pSNW2-PP_5388-*P*_*14c*_*-BCD22-phi15lys(G3RQ)*, which integrated in the *P. putida* KT2440 and *P. putida* SEM11 hosts after electroporation. After overnight incubation of several transformants, the pSEVA62313S helper vector with constitutive expression of I-SceI was introduced into the hosts by electroporation. As recommended in the original protocol (36), the resulting colonies were transferred to a fresh LB agar plate by streaking to avoid mixed-phenotype colonies. The resulting colonies were screened for a successful second HR event by verifying the lack of green fluorescence, followed by a PCR with the same genomic primers as for the CRISPR-Cas3-based method. For *P. putida* KT2440, only one of 24 PCR-screened colonies contained a correct integrant, while for *P. putida* SEM11 no correct integrants were obtained (0/24), but still appeared to have a mixed phenotype (Figure 3a, Figure S3). Therefore, the *P. putida* SEM11 strain carrying pSNW2-PP_5388-*P*_*14c*_*-BCD22-phi15lys(G3RQ)* and pSEVA62313S helper vector was streaked twice more to allow additional time for the second HR event to occur. After a second PCR screen, 21% (5/24) of the screened colonies showed an amplicon length correlating to correct integration of the *P*_*14c*_*-BCD22-phi15lys(G3RQ)* cassette (Figure 3a, Figure S3). Overall, the engineering efficiencies obtained by homologous recombination were much lower compared to the CRISPR-Cas3-based method and required significantly more handling time, due to consecutive electroporation of the vectors and multiple streaking steps.

### The CRISPR-Cas3 system cures itself with high efficiency in *P. putida* using an *oriT*-targeting spacer

After successful engineering of the host genome, cells need to be cured from the pCas3cRh and repair template vector for downstream processing. A universal CRISPR-Cas3-based curing concept was introduced, similar to the proven CRISPR-Cas9-based curing method for *E. coli* and *P. putida*, which makes use of spacers targeting conserved regions of plasmid, i.e. the origins-of-replication (*oriR*) (42). As the cells in this work already contained the Cas3 system, it can simply be used to target itself by introducing crRNA with a self-targeting spacer. To this end, a universal spacer was designed, binding specifically to the *oriT* located on all SEVA plasmids and many other commonly used vectors for genome engineering, including pCas3cRh. This *oriT* spacer and crRNA were cloned into pSEVA521 under control of the *P*_*RhaBAD*_ promoter (Tables S1 and S2) and called pSEVA52-oriT.

The pSEVA52-oriT vector was introduced in the engineered *P. putida* KT-phi15lys and *P. putida* S-phi15lys strains through electroporation, after which cells were plated on LB agar supplemented with gentamicin (pCas3cRh-PP_5388) and tetracycline (pSEVA52-oriT). In parallel, the same strains were electroporated with pSEVA521 as a negative control. The following day, several colonies of each strain were grown in LB^Gm10/Tc10^ medium with 0.1% rhamnose to induce expression of the Cas3 system. After a dilution streak on LB medium without any antibiotics and overnight incubation, 24 colonies of each condition were screened on gentamicin (pCas3cRh-PP_5388), kanamycin (pSEVA23-PP_5388) and tetracycline (pSEVA52-oriT) to assess the curing efficiency. In the presence of the *oriT* spacer, 91.6% and 66.7% of colonies were fully cured of all vectors for *P. putida* KT2440 and *P. putida* SEM11, respectively (Figure 3b). This is in sharp contrast to the control samples with the empty pSEVA521 vector, of which all of the screened colonies still contained at least two of the three vectors. Furthermore, the engineered strains were also subjected to serial passaging for the same amount of time as required for the CRISPR-Cas3-based curing. Four passages were performed over three days, after which none of the screened colonies were cured from the pCas3cRh and pSEVA23-PP_5388 vectors (Figure 3b). These results show that the CRISPR-Cas3-system is able to efficiently target itself and other vectors in the same cell, with enhanced efficiencies compared to the original CRISPR-Cas9-based curing approach (53% curing efficiency in *P. putida*) (42).

After successful vector curing, two biological replicates of *P. putida* KT-phi15lys and *P. putida* S-phi15lys were subjected to whole genome sequencing. No substantial deletions or insertions were detected, except for four point mutations outside of the integrated region in the *P. putida* KT-phi15lys replicates (Tables S5 and S6) and two point mutations in both *P. putida* S-phi15lys replicates (Tables S7 and S8).

### Application examples: efficient genomic deletion of three different targets in *P. aeruginosa*

To show that the CRISPR-Cas3-based engineering method is also functional in other hosts, three separate genomic deletions were created in the genome of *P. aeruginosa* PAO1. More specifically, three sets of spacers and repair templates were designed to delete the entire coding sequences of *fleS*, PA_2560 and *prpL* (Figure 4a). After successful construction of all six vectors, the corresponding pCas3cRh-spacer and pSEVA13-HDR vectors were simultaneously introduced in *P. aeruginosa* PAO1. Surprisingly, visible colonies only appeared after a two-day incubation period for PA_2560 and *prpL*, while a control electroporation with the empty pCas3cRh or pSEVA131 vector resulted in colony formation overnight. For the *fleS* deletion, even after multiple days of incubation, no colonies grew on plates of the co-electroporation and the pSEVA13-HDR and pCas3cRh vectors had to be introduced consecutively. This is in sharp contrast to the results with *P. putida*, where co-electroporation resulted in normal colony formation after a single night of incubation.

**Figure 4:**
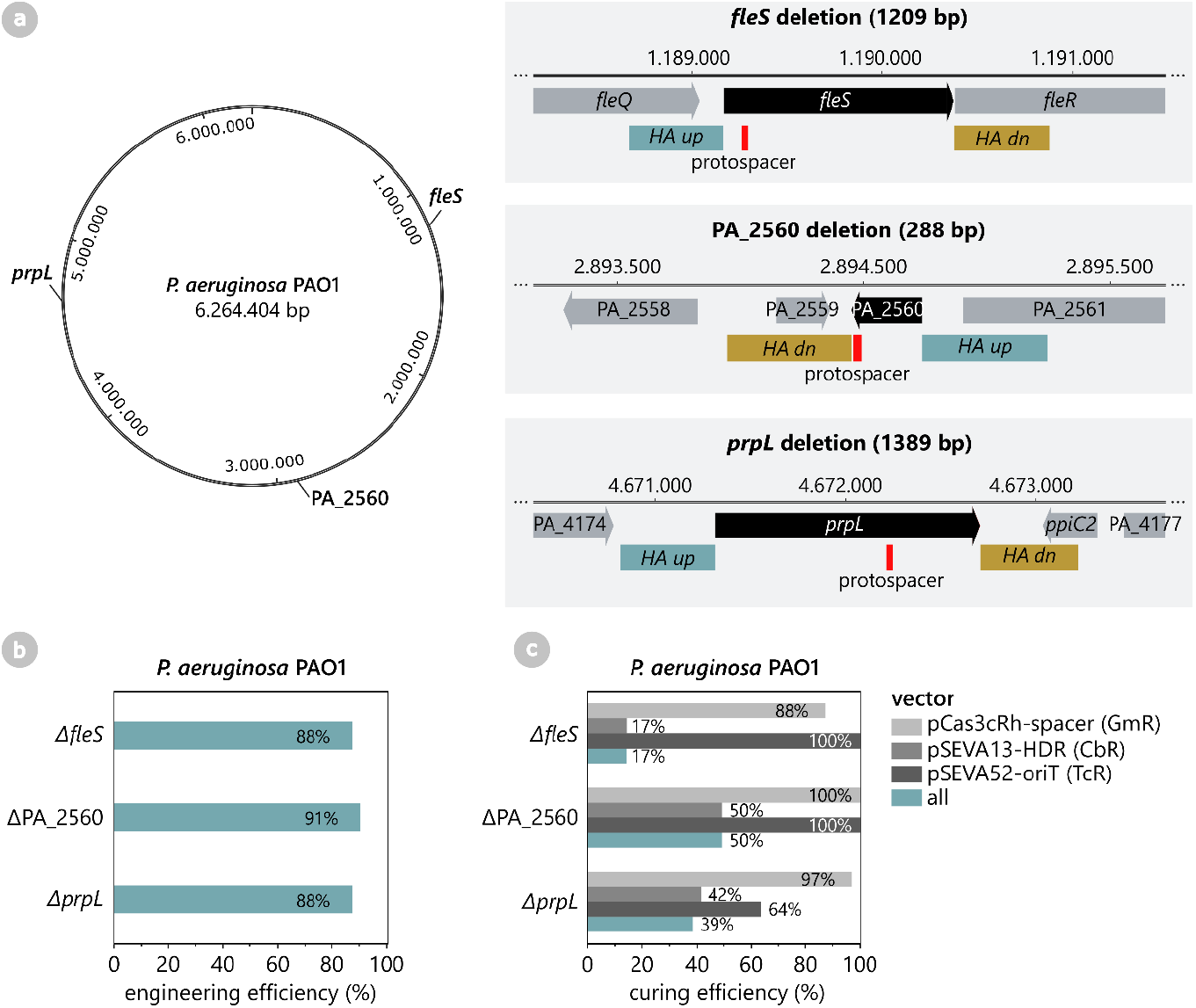
a) Three separate genomic deletions are created in the *P. aeruginosa* PAO1 genome, namely the entire coding regions of *fleS*, PA_2560 and *prpL* (indicated in black). The position of the protospacer in the genes is marked in red, and the upstream and downstream homology arm are indicated in cyan and ochre, respectively. b) Engineering efficiency of the CRISPR-Cas3-based engineering method to create the *fleS*, PA_2560 and *prpL* deletions. c) Curing efficiency of the CRISPR-Cas3-based curing method for pCas3cRh-spacer, pSEVA13-HDR and pSEVA52-oriT in the *P. aeruginosa* PAO1 *ΔfleS*, ΔPA_2560 and *ΔprpL* deletion mutants.

This indicates that *P. aeruginosa* shows retarded cell growth after introducing the engineering vectors, which points towards significant basal Cas3 expression from the RhaRS/*P*_*rhaBAD*_ system in this host. To confirm this hypothesis, the CRISPR-Cas3 system was not induced with rhamnose, but the transformants were directly after electroporation analyzed with PCR for the genomic deletion. Indeed, for all three deletions, at least 88% of the screened colonies already contained the desired deletion (Figure 4b, Figures S4-6). This confirms that in *P. aeruginosa*, the basal expression of the CRISPR-Cas3-system is sufficient for genome engineering and rhamnose induction is not required. Other inducible systems could be explored to create a more stringent regulation of the Cas3 system in *P. aeruginosa*.

After successful deletion of the targeted genes, all three deletion mutants were cured from the respective pCas3cRh-spacer and pSEVA13-HDR vectors using the *oriT*-targeting approach. The pSEVA52-oriT vector was introduced in all strains, after which the transformants were grown overnight in antibiotic-free medium without rhamnose. The following day, a dilution streak was performed and the resulting colonies were screened for sensitivity against gentamycin (pCas3cRh-spacer), carbenicillin (pSEVA13-HDR) and tetracycline (pSEVA52-oriT). Both the pCas3cRh-spacer and pSEVA52-oriT vectors were cured very effectively, with a curing efficiency ranging from 64 to 100% (Figure 4c). The pSEVA13-HDR vectors, on the other hand, were still present in the majority of the screened colonies, resulting in a rather low curing efficiency of 17%, 42% and 50% for the Δ*fleS*, Δ*prpL* and ΔPA_2560 mutants, respectively. This difference in curing efficiency between the vectors could be explained by the fact that the pCas3cRh vector exerts a negative selection pressure upon itself once pSEVA52-oriT is present, in contrast to the other two vectors. Furthermore, the pSEVA52-oriT vector contains the low-copy RK2 *oriR*, while the pSEVA13-HDR vector carries the medium-copy BBR1 *oriR*, which could explain why the pSEVA52-oriT origin is more efficiently cured than its pSEVA13-HDR counterpart. To further improve the flexibility of the system and the efficiencies achieved, additional spacers could be included on pSEVA52-oriT to target a variety of *oriRs*, shown to be effective in previous work (42).

### An easy-to-clone vector set with a broad range of antibiotic markers further improves the CRISPR-Cas3-based engineering method

Two vector sets were created to facilitate cloning of the homology arms and to allow compatibility of the CRISPR-Cas3 engineering system with different hosts or experimental set-ups requiring different antibiotic selection markers. The first vector set for HR cloning comprises five pSEVAX3-GG vectors, all encoding a Golden Gate cassette and different antibiotic markers (Figure 5a). The Golden Gate cassette consists of an msfGFP (<u>m</u>onomeric <u>s</u>uperfolder <u>g</u>reen fluorescent <u>p</u>rotein) reporter driven by a strong constitutive promoter (*P*_*14g*_) (48) and flanking BsaI recognition sites (Figure 5c). The second vector set, on the other hand, is derived from pCas3cRh and holds five pCas3-Ab vectors with different antibiotic markers (Figure 5b). As such, the user has the possibility to select their favorite vector combination for the genomic engineering experiment in mind.

**Figure 5:**
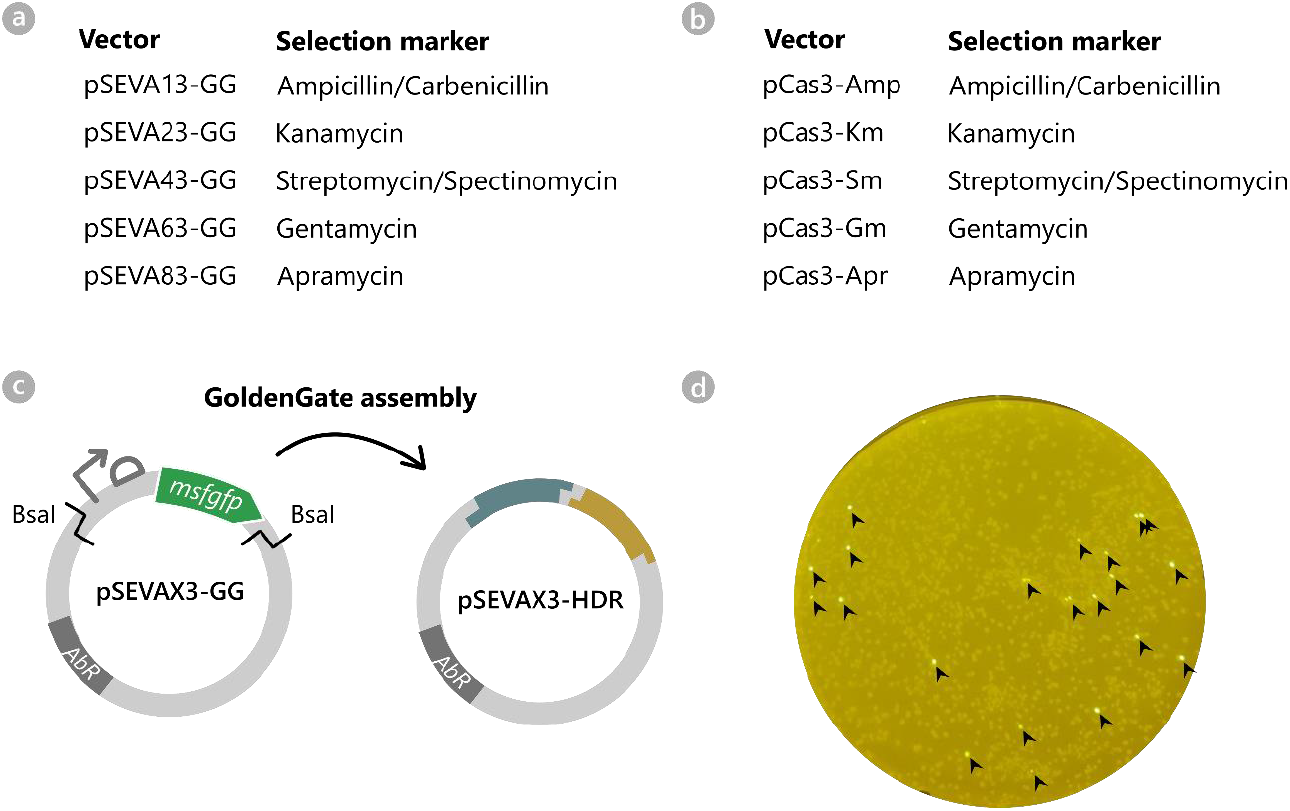
a) Vector set of pSEVAX3-GG for Golden Gate cloning of the homology arms for CRISPR-Cas3 engineering. All vectors are identical, except for the antibiotic selection marker. The vectors are equipped with a Golden Gate cassette, consisting of an msfGFP reporter flanked with BsaI recognition sites. b) Vector set of pCas3cRh-derived vectors for CRISPR-Cas3-based genome engineering. All vectors are identical, except for the antibiotic selection marker. c) Golden Gate assembly of the homology arms into pSEVAX3-GG vectors. The Golden Gate cassette with msfGFP reporter is substituted for the homology arms for HDR (teal and yellow). d) LB agar plate on an ultrabright-LED transilluminator (470 nm): *E. coli* after transformation with assembled pSEVAX3-HDR vector (Golden Gate reaction mix). Colonies which are false-positive and thus contain the original pSEVAX3-GG vector are easily identified by their msfGFP fluorescence (indicated with a black arrow).

### Conclusions and Perspectives

A novel CRISPR-Cas3-assisted editing method was presented for *Pseudomonas*, showcasing high efficiency for genomic integration or deletion in *P. putida* and *P. aeruginosa* (>83%). In addition, due to the inherent ability of the Cas3 enzyme to cleave and thereby cure plasmids from the host strain, all helper vectors are rapidly and effectively removed in only a few days, with up to 100% curing efficiency. As such, the described approach is an elegant addition to the CRISPR-Cas-based engineering toolbox for *Pseudomonas*. Apart from their use in genomic engineering, the pCas3-AbR and pSEVA52-oriT vectors can be used as a stand-alone tool for vector curing of any synthetic or naturally-occurring plasmid in *Pseudomonas*. By integrating the *oriT* spacer on the pCas3-AbR vector under control of a strictly regulated promoter, only a single vector would be required for curing purposes.

In future work, the possibilities of the Cas3 editing approach can be further explored, e.g. by creating larger genomic alterations or by performing several genomic edits simultaneously, by providing more than one spacer and repair template on the pCas3-AbR and pSEVAX3-GG plasmids. Additionally, due to the flexibility of the proposed vectors sets, namely the pSEVAX3-GG and pCas3-AbR sets, the functionality of the Cas3 approach can readily be investigated in related *Pseudomonas* species or other Gram-negative strains.

## Materials and Methods

### Strains and Media

All strains used in this work are listed in Table S2. Overall, vector construction was performed in *E. coli* TOP10 and CRISPR-Cas3-based engineering was carried out in *P. putida* KT2440, *P. putida* SEM11 and *P. aeruginosa* PAO1 (Table S2). All strains were cultured in standard LB medium or agar, supplemented with the appropriate antibiotics: Gm10 (*E. coli* and *P. putida*) or Gm50 (*P. aeruginosa*), Km50 (*E. coli* and *P. putida*), Ap100 (*E. coli*), Cb200 (*P. aeruginosa*), Tc10 (*E. coli* and *P. putida*) or Tc60 (*P. aeruginosa*). *P. putida* was incubated at 30°C, whereas *E. coli* and *P. aeruginosa* were incubated at 37°C.

### Vector construction – pCas3cRh-spacer

A spacer sequence was identified in the target region and introduced in the pCas3cRh vector by Golden Gate ligation. First, the spacer was created by annealing two primers: 1) GAAAC-[spacer sequence]-G and 2) GCGAC-[reverse complement of spacer sequence]-G. The primers used in this work are listed in Table S1. The annealed primer pair (50 ng) was combined with pCas3cRh (100 ng), T4 DNA ligase (1 U, Thermo Scientific), BsaI (10 U, Thermo Scientific) and 1x DNA ligation buffer (Thermo Scientific), after which the reaction mixture was subjected to 30 restriction-ligation cycles (37°C for 2 min; 16°C for 3 min). Next, the reaction mixture was introduced in *E. coli* TOP10 via heat-shock transformation (49). After overnight incubation on LB^Gm10^ agar, multiple transformants were screened for the presence of the spacer using DreamTaq Green PCR (Thermo Scientific) with primers pCas3cRh_F/R (Table S1). Amplicons with the expected length were Sanger sequenced (Eurofins Genomics, Germany) and corresponding vectors were purified with the GeneJet Miniprep Kit (Thermo Scientific) (Table S2).

### Vector construction – pSEVAX3-HDR

The template for HDR is provided on pSEVA131 (*P. aeruginosa*) or pSEVA231 (*P. putida*), further referred as pSEVAX31, and assembled by Golden Gate cloning. First, the upstream and downstream homology arms (HA up and dn), desired insert (for integrations only) and the vector backbone were amplified with Phusion polymerase (Thermo Scientific) with tailed primers, to introduced the BsaI recognition site and BsaI restriction site for Golden Gate ligation (Table S1). All nucleotide sequences of used HAs and inserts in this work are provided in Table S4. The BsaI restriction sites are designed to allow specific annealing of HA up – (insert) – HA dn in the pSEVAX3 amplicon. The amplicons of the homology arms (50 ng) each and insert (50 ng) were combined with linearized pSEVAX3 (100 ng), T4 DNA ligase (1 U, Thermo Scientific), BsaI (10 U, Thermo Scientific) and 1x DNA ligation buffer (Thermo Scientific), after which the reaction mixture was subjected to 50 restriction-ligation cycles (37°C for 2 min; 16°C for 3 min). Next, the reaction mixture was introduced in *E. coli* TOP10 via heat-shock transformation (49). After overnight incubation on LB^Km50^ or LB^Ap100^ agar, multiple transformants were screened for the presence of the template using DreamTaq Green PCR (Thermo Scientific) with primers SEVA_PS1/2 (Table S1). Amplicons of the expected length were Sanger sequenced (Eurofins Genomics, Germany) and corresponding vectors were purified with the GeneJet Miniprep Kit (Thermo Scientific) (Table S2).

### Vector construction – pSEVA52-oriT

Vector pSEVA52-oriT was constructed in two steps. First, a pCas3cRh vector with *oriT* spacer was constructed as described above, with oriT_spacer_F/R (Tables S1 and S2). Second, the *P*_*RhaBAD*_ promoter and CRISPR array with *oriT* spacer were amplified from pCas3cRh-oriT with tailed primers oriT_Cas3_F/R and the pSEVA521 backbone was linearized with Phusion PCR (Thermo Scientific) with tailed primers oriTcas3_SEVA_F/R (Table S1). Both amplicons were annealed by Golden Gate ligation, as described above for the construction of pSEVAX3-HDR vectors. Multiple *E. coli* TOP10 transformants were screened for the presence of the *oriT* CRISPR array using DreamTaq Green PCR (Thermo Scientific) with primers SEVA_PS1/2 (Table S1). Amplicons of the expected length were Sanger sequenced (Eurofins Genomics, Germany) and the final pSEVA52-oriT vector was purified with the GeneJet Miniprep Kit (Thermo Scientific) (Table S2).

### Vector construction – pCas3-XX and pSEVAX3-GG vector sets

To create Cas3 bearing plasmids with different antibiotic selection markers (pCas3-Amp, pCas3-Km, pCas3-Sm, pCas3-Gm and pCas3-Apr; Table S2), pCas3cRh was amplified with primer pair pCas3_Ab_F/R (Table S1) and the antibiotic selection cassettes were amplified from canonical SEVA plasmids (44) with the primer pair Ab_F/R. The antibiotic selection fragments were ligated with the pCas3cRh amplicon by USER cloning (50). Following transformation of *E. coli*, colony PCR and plasmid purification as described above, correctness of plasmids was confirmed by whole plasmid sequencing (Plasmidsaurus, Oregon, USA).

For the creation of the pSEVAX3-GG vector set, pSEVA131 was amplified with the primer pair pSX31_GG_F/R, while a fragment carrying *msfgfp* under the constitutive promoter 14g with BCD2 was amplified from pBG42 (48) with primer pair P14g-BCD2-GFP_F/R. Fragments were merged by USER cloning into pSEVA13-GG and correctness of the plasmid inserts was confirmed by Sanger sequencing with SEVA_PS1/2. The overhangs created by BsaI were designed for optimal cloning efficiency (51). Subsequently, the antibiotic cassette of the plasmid was exchanged by USER cloning to create pSEVA23-GG, pSEVA43-GG, pSEVA63-GG and pSEVA83-GG (Table S2). The vector, linearized with the primer pair pSX31_Ab_F/R, was merged with the same fragments used for antibiotic cassette exchange for pCas3cRh. Correct vector assembly was verified with nanopore, whole plasmid sequencing. Finally, vectors pCas3-ApR and pSEVA83-GG were subjected to full linearization with a tailed primer (ApR_BsaI_F/R) and religated with USER cloning, to remove an undesired BsaI recognition site from the *apR* gene.

### Electroporation

*P. putida* and *P. aeruginosa* were electroporated according to the protocol described by Choi *et al*. (52). In brief, overnight cultures were washed three to five times in a sterile 10% sucrose solution to create electrocompetent cells. After the washing steps, 20-50 ng plasmid DNA was added to a 100 μL cell aliquot and electroshocked at 200 ohm, 25 μF, and 1.8 kV or 2.0 kV for *P. aeruginosa* and *P. putida*, respectively. For co-electroporations, 100 ng of each plasmid was added to the cell aliquot together and electroshocked in the same manner. After cell recovery for 1.5h in LB or SOC medium at the appropriate temperature, cells were plated on selective LB agar and incubated overnight, unless specifically mentioned otherwise.

### CRISPR-Cas3-based engineering and vector curing in *P. putida*

Overnight cultures of *P. putida* were co-electroporated with pCas3cRh-PP_5388 and pSEVA23-PP_5388 as described above. After overnight incubation on LBKm50/Gm10 agar, five colonies were inoculated together in 20 mL LB^Km50/Gm10^ with 0.1% rhamnose (Merck, CAS no. 10030-85-0) for induction of the CRISPR-Cas3 system and incubated overnight while shaking. The next day, a dilution streak of the 20 mL culture is performed on LB^Km50/Gm10^ agar and again incubated overnight, after which 24 colonies were screened for correct genomic integration of the insert with DreamTaq Green PCR (Thermo Scientific) with primers PP5388_up/dn (Table S1). Amplicons of the expected length were Sanger sequenced (Eurofins Genomics, Germany) and the corresponding colonies were cured from pCas3cRh-PP_5388 and pSEVA23-PP_5388. For vector curing, overnight cultures were electroporated with pSEVA521-oriT and the CRISPR-Cas3 system is induced as mentioned previously, using overnight incubation with 0.1% rhamnose followed by a dilution streak on LB medium without antibiotics. From the resulting plates, 24 colonies were streaked on LB, LB^Km50^, LB^Gm10^ and LB^Tc10^ and incubated overnight to assess successful vector curing by antibiotic sensitivity.

### CRISPR-Cas3-based engineering and vector curing in *P. aeruginosa*

Overnight cultures of *P. aeruginosa* were co-electroporated with pCas3cRh-spacer and pSEVA131-HDR as described above. For deletion of *fleS*, the co-electroporation did not result in colony formation, such that pSEVA13-FleS and pCas3cRh-FleS were introduced consecutively. After a two-day incubation period on LB^Cb200/Gm10^ agar, 14-24 colonies were screened for correct genomic deletion of the target gene with DreamTaq Green PCR (Thermo Scientific) with primers gene_up/dn (Table S1). Amplicons of the expected length were Sanger sequenced (Eurofins Genomics, Germany) and the corresponding colonies were cured from pCas3cRh-spacer and pSEVA13-HDR. For vector curing, overnight cultures were electroporated with pSEVA52-oriT and incubated overnight. The following day, 24 colonies were streaked on LB, LB^Cb200^, LB^Gm50^ and LB^Tc60^ and incubated overnight to assess successful vector curing by antibiotic sensitivity.

### Whole-genome sequencing

The genomic DNA of the CRISPR-Cas3 engineered strains after vector curing was isolated using the DNeasy UltraClean Microbial Kit (Qiagen, Germany) according to the manufacturer’s guidelines. The obtained DNA was sequenced with an Illumina platform (USA) and an Oxford Nanopore Technologies platform (UK) for long-read DNA sequencing. The Illumina DNA libraries were prepared using the Illumina DNA Prep kit (USA) and the Nextera™ DNA CD Indexes (Illumina, USA). The average length of the DNA libraries was evaluated using Agilent Bioanalyzer 2100 and a High Sensitivity Kit (Agilent Technologies, USA) and the concentration of the DNA libraries was determined with a Qubit device (Thermo Fisher Scientific, USA). Next, the samples were pooled together for sequencing on the Illumina MiniSeq NGS platform. The MiniSeq Mid Output Kit (300-cycles) (Illumina, USA) was used for paired-end sequencing (2x150 bp), aiming for 800 000 reads per sample.

For Nanopore sequencing, the Rapid Barcoding Kit 24 V14 (Oxford Nanopore Technologies, UK) was used for library preparation. A maximum of 24 samples were pooled and sequenced on a R10.4.1 flowcell (Oxford Nanopore Technologies, UK). The raw Illumina and Nanopore reads were trimmed with Trimmomatic (53) or Porechop (54), respectively, after which they were assembled into complete circular genomes with Unicycler (55). Large deletions were visualized in IGV after Bowtie2 assembly (56) and SNP analysis was performed with SNIPPY (57).

## Data availability

All essential data supporting this article is provided in the main text or the supporting information.

## Acknowledgements

The pCas3cRh vector was kindly provided by prof. J. Bondy-Denomy (UCSF). This project received funding from the European Research Council (ERC) under the European Union’s Horizon 2020 Research and Innovation Programme (Grant Agreements 819800 and 814418), from the Fonds voor Wetenschappelijk Onderzoek Vlaanderen (FWO) as part of the CELL-PHACTORY Project (Grant G096519N), from the Novo Nordisk Foundation (Grant Agreements NNF10CC1016517 and NNF18CC0033664) and by a grant from KU Leuven (C1 project ‘ACES’, C16/20/001).

## References

1. Garrity GM, Bell JA, Lilburn T. 2005. Pseudomonadales Orla-Jensen 1921, 270AL, p. 323–442. In Brenner, DJ, Krieg, NR, Staley, JT, Garrity, GM, Boone, DR, De Vos, P, Goodfellow, M, Rainey, FA, Schleifer, K-H (eds.), Bergey’s Manual® of Systematic Bacteriology: Volume Two The Proteobacteria Part B The Gammaproteobacteria. Springer US, Boston, MA.

2. Breidenstein EBM, de la Fuente-Nunez C, Hancock REW. 2011. Pseudomonas aeruginosa: all roads lead to resistance. Trends Microbiol 19:419–426.

3. Nikel PI, Chavarría M, Danchin A, de Lorenzo V. 2016. From dirt to industrial applications: Pseudomonas putida as a Synthetic Biology chassis for hosting harsh biochemical reactions. Curr Opin Chem Biol 34:20–29.

4. Nikel PI, de Lorenzo V. 2018. Pseudomonas putida as a functional chassis for industrial biocatalysis: From native biochemistry to trans-metabolism. Metab Eng 50:142–155.

5. Lister PD, Wolter DJ, Hanson ND. 2009. Antibacterial-resistant Pseudomonas aeruginosa: clinical impact and complex regulation of chromosomally encoded resistance mechanisms. Clin Microbiol Rev 22:582–610.

6. Choi KH, Gaynor JB, White KG, Lopez C, Bosio CM, Karkhoff-Schweizer RAR, Schweizer HP. 2005. A Tn7-based broad-range bacterial cloning and expression system. Nat Methods 2:443– 448.

7. Weihmann R, Domröse A, Drepper T, Jaeger K-E, Loeschcke A. 2020. Protocols for yTREX/Tn5-based gene cluster expression in Pseudomonas putida. Microb Biotechnol 13:250–262.

8. Hmelo LR, Borlee BR, Almblad H, Love ME, Randall TE, Tseng BS, Lin C, Irie Y, Storek KM, Yang JJ, Siehnel RJ, Howell PL, Singh PK, Tolker-Nielsen T, Parsek MR, Schweizer HP, Harrison JJ. 2015. Precision-engineering the Pseudomonas aeruginosa genome with two-step allelic exchange. Nat Protoc 10:1820–1841.

9. Martínez-García E, de Lorenzo V. 2011. Engineering multiple genomic deletions in Gram-negative bacteria: Analysis of the multi-resistant antibiotic profile of Pseudomonas putida KT2440. Environ Microbiol 13:2702–2716.

10. Liang R, Liu J. 2010. Scarless and sequential gene modification in Pseudomonas using PCR product flanked by short homology regions. BMC Microbiol 10:209.

11. Chen Z, Ling W, Shang G. 2016. Recombineering and I-SceI-mediated Pseudomonas putida KT2440 scarless gene deletion. FEMS Microbiol Lett 363:1–7.

12. Luo X, Yang Y, Ling W, Zhuang H, Li Q, Shang G. 2016. Pseudomonas putida KT2440 markerless gene deletion using a combination of λ Red recombineering and Cre/loxP site-specific recombination. FEMS Microbiol Lett 363.

13. Choi KR, Cho JS, Cho IJ, Park D, Lee SY. 2018. Markerless gene knockout and integration to express heterologous biosynthetic gene clusters in Pseudomonas putida. Metab Eng 47:463– 474.

14. Jiang W, Bikard D, Cox D, Zhang F, Marraffini LA. 2013. RNA-guided editing of bacterial genomes using CRISPR-Cas systems. Nat Biotechnol 31:233–239.

15. McAllister KN, Bouillaut L, Kahn JN, Self WT, Sorg JA. 2017. Using CRISPR-Cas9-mediated genome editing to generate C. difficile mutants defective in selenoproteins synthesis. Sci Rep 7:14672.

16. Oh J-H, van Pijkeren J-P. 2014. CRISPR-Cas9-assisted recombineering in Lactobacillus reuteri. Nucleic Acids Res 42:e131.

17. DiCarlo JE, Norville JE, Mali P, Rios X, Aach J, Church GM. 2013. Genome engineering in Saccharomyces cerevisiae using CRISPR-Cas systems. Nucleic Acids Res 41:4336–4343.

18. Jiang W, Zhou H, Bi H, Fromm M, Yang B, Weeks DP. 2013. Demonstration of CRISPR/Cas9/sgRNA-mediated targeted gene modification in Arabidopsis, tobacco, sorghum and rice. Nucleic Acids Res 41:e188.

19. Mali P, Yang L, Esvelt KM, Aach J, Guell M, DiCarlo JE, Norville JE, Church GM. 2013. RNA-guided human genome engineering via Cas9. Science 339:823–826.

20. Shuman S, Glickman MS. 2007. Bacterial DNA repair by non-homologous end joining. Nat Rev Microbiol 5:852–861.

21. Wirth NT, Kozaeva E, Nikel PI. 2019. Accelerated genome engineering of Pseudomonas putida by I-SceI–mediated recombination and CRISPR-Cas9 counterselection. Microb Biotechnol 13:233–249.

22. Aparicio T, de Lorenzo V, Martínez-García E. 2018. CRISPR/Cas9-Based Counterselection Boosts Recombineering Efficiency in Pseudomonas putida. Biotechnol J 13:e1700161.

23. Sun J, Wang Q, Jiang Y, Wen Z, Yang L, Wu J, Yang S. 2018. Genome editing and transcriptional repression in Pseudomonas putida KT2440 via the type II CRISPR system. Microb Cell Fact 17:1–17.

24. Chen W, Zhang Y, Zhang Y, Pi Y, Gu T, Song L, Wang Y, Ji Q. 2018. CRISPR/Cas9-based Genome Editing in Pseudomonas aeruginosa and Cytidine Deaminase-Mediated Base Editing in Pseudomonas Species. iScience 6:222–231.

25. Wu Z, Chen Z, Gao X, Li J, Shang G. 2019. Combination of ssDNA recombineering and CRISPR-Cas9 for Pseudomonas putida KT2440 genome editing. Appl Microbiol Biotechnol 103:2783– 2795.

26. Zhou Y, Lin L, Wang H, Zhang Z, Zhou J, Jiao N. 2020. Development of a CRISPR/Cas9n-based tool for metabolic engineering of Pseudomonas putida for ferulic acid-to-polyhydroxyalkanoate bioconversion. Commun Biol 3:1–13.

27. Cook TB, Rand JM, Nurani W, Courtney DK, Liu SA, Pfleger BF. 2018. Genetic tools for reliable gene expression and recombineering in Pseudomonas putida. J Ind Microbiol Biotechnol 45:517–527.

28. Wirth NT, Kozaeva E, Nikel PI. 2020. Accelerated genome engineering of Pseudomonas putida by I-SceI-mediated recombination and CRISPR-Cas9 counterselection. Microb Biotechnol 13:233–249.

29. Lin Z, Li H, He L, Jing Y, Pistolozzi M, Wang T, Ye Y. 2021. Efficient genome editing for Pseudomonas aeruginosa using CRISPR-Cas12a. Gene 790:145693.

30. Xu Z, Li M, Li Y, Cao H, Miao L, Xu Z, Higuchi Y, Yamasaki S, Nishino K, Woo PCY, Xiang H Yan 2019. Native CRISPR-Cas-Mediated Genome Editing Enables Dissecting and Sensitizing Clinical Multidrug-Resistant P. aeruginosa. Cell Rep 29:1707–1717.e3.

31. Csörgő B, León LM, Chau-Ly IJ, Vasquez-Rifo A, Berry JD, Mahendra C, Crawford ED, Lewis JD, Bondy-Denomy J. 2020. A compact Cascade-Cas3 system for targeted genome engineering. Nat Methods 17:1183–1190.

32. Zimmermann A, Verstrepen KJ, Prieto-vivas JE, Cautereels C. 2023. A Cas3-base editing tool for targetable in vivo mutagenesis https://doi.org/10.1038/s41467-023-39087-z.

33. Trevors JT. 1986. Plasmid curing in bacteria. FEMS Microbiol Rev 1:149–157.

34. Martínez-García E, Aparicio T, de Lorenzo V, Nikel PI. 2017. Engineering Gram-Negative Microbial Cell Factories Using Transposon Vectors. Methods Mol Biol 1498:273–293.

35. Reyrat JM, Pelicic V, Gicquel B, Rappuoli R. 1998. Counterselectable markers: untapped tools for bacterial genetics and pathogenesis. Infect Immun 66:4011–4017.

36. Volke DC, Wirth NT, Nikel PI. 2021. Rapid Genome Engineering of Pseudomonas Assisted by Fluorescent Markers and Tractable Curing of Plasmids. bio-protocol 11:1–16.

37. Volke DC, Friis L, Wirth NT, Turlin J, Nikel PI. 2020. Synthetic control of plasmid replication enables target- and self-curing of vectors and expedites genome engineering of Pseudomonas putida. Metab Eng Commun 10:e00126.

38. Crameri R, Davies JE, Hütter R. 1986. Plasmid curing and generation of mutations induced with ethidium bromide in streptomycetes. J Gen Microbiol 132:819–824.

39. Jäger W, Schäfer A, Pühler A, Labes G, Wohlleben W. 1992. Expression of the Bacillus subtilis sacB gene leads to sucrose sensitivity in the gram-positive bacterium Corynebacterium glutamicum but not in Streptomyces lividans. J Bacteriol 174:5462–5465.

40. Chen S, Larsson M, Robinson RC, Chen SL. 2017. Direct and convenient measurement of plasmid stability in lab and clinical isolates of E. coli. Sci Rep 7:4788.

41. Karunakaran P, Blatny JM, Ertesvåg H, Valla S. 1998. Species-dependent phenotypes of replication-temperature-sensitive trfA mutants of plasmid RK2: a codon-neutral base substitution stimulates temperature sensitivity by leading to reduced levels of trfA expression. J Bacteriol 180:3793–3798.

42. Lauritsen I, Porse A, Sommer MOA, Nørholm MHH. 2017. A versatile one-step CRISPR-Cas9 based approach to plasmid-curing. Microb Cell Fact 16:1–10.

43. Lauritsen I, Kim SH, Porse A, Nørholm MHH. 2018. Standardized Cloning and Curing of Plasmids. Methods Mol Biol 1772:469–476.

44. Martínez-García E, Fraile S, Algar E, Aparicio T, Velázquez E, Calles B, Tas H, Blázquez B, Martín B, Prieto C, Sánchez-Sampedro L, Nørholm MHH, Volke DC, Wirth NT, Dvořák P, Alejaldre L, Grozinger L, Crowther M, Goñi-Moreno A, Nikel PI, Nogales J, de Lorenzo V. 2023. SEVA 4.0: an update of the Standard European Vector Architecture database for advanced analysis and programming of bacterial phenotypes. Nucleic Acids Res 51:D1558–D1567.

45. Lammens E-M, Feyaerts N, Kerremans A, Boon M, Lavigne R. 2023. Assessing the Orthogonality of Phage-Encoded RNA Polymerases for Tailored Synthetic Biology Applications in Pseudomonas Species. Int J Mol Sci 24:1–18.

46. Chaves JE, Wilton R, Gao Y, Munoz NM, Burnet MC, Schmitz Z, Rowan J, Burdick LH, Elmore J, Guss A, Close D, Magnuson JK, Burnum-Johnson KE, Michener JK. 2020. Evaluation of chromosomal insertion loci in the Pseudomonas putida KT2440 genome for predictable biosystems design. Metab Eng Commun 11:e00139.

47. Martin-Pascual M, Batianis C, Bruinsma L, Asin-Garcia E, Garcia-Morales L, Weusthuis RA, van Kranenburg R, Martins dos Santos VAP. 2021. A navigation guide of synthetic biology tools for Pseudomonas putida. Biotechnol Adv 49:107732.

48. Zobel S, Benedetti I, Eisenbach L, De Lorenzo V, Wierckx N, Blank LM. 2015. Tn7-based device for calibrated heterologous gene expression in Pseudomonas putida. ACS Synth Biol 4:1341– 1351.

49. Green R, Rogers EJ. 2013. Transformation of chemically competent E. coli, p. 329–336. In Methods in Enzymology, 1st ed. Elsevier Inc.

50. Frandsen RJN, Andersson JA, Kristensen MB, Giese H. 2008. Efficient four fragment cloning for the construction of vectors for targeted gene replacement in filamentous fungi. BMC Mol Biol 9:1–11.

51. Pryor JM, Potapov V, Kucera RB, Bilotti K, Cantor EJ, Lohman GJS. 2020. Enabling one-pot Golden Gate assemblies of unprecedented complexity using data-optimized assembly design. PLoS One 15:1–19.

52. Choi KH, Kumar A, Schweizer HP. 2006. A 10-min method for preparation of highly electrocompetent Pseudomonas aeruginosa cells: Application for DNA fragment transfer between chromosomes and plasmid transformation. J Microbiol Methods 64:391–397.

53. Bolger AM, Lohse M, Usadel B. 2014. Trimmomatic: A flexible trimmer for Illumina sequence data. Bioinformatics 30:2114–2120.

54. Becker K, Meyer A, Roberts TM, Panke S. 2021. Plasmid replication based on the T7 origin of replication requires a T7 RNAP variant and inactivation of ribonuclease H. Nucleic Acids Res 49:8189–8198.

55. Wick RR, Judd LM, Gorrie CL, Holt KE. 2017. Unicycler: Resolving bacterial genome assemblies from short and long sequencing reads. PLoS Comput Biol 13:1–22.

56. Langmead B, Salzberg SL. 2012. Fast gapped-read alignment with Bowtie 2. Nat Methods 9:357–359.

57. Seemann T. 2015. SNIPPY: fast bacterial variant calling from NGS reads.

58. Csörgő B, León LM, Chau-Ly IJ, Vasquez-Rifo A, Berry JD, Mahendra C, Crawford ED, Lewis JD, Bondy-Denomy J. 2020. A compact Cascade–Cas3 system for targeted genome engineering. Nat Methods 17:1183–1190.

59. Silva-Rocha R, Martínez-García E, Calles B, Chavarría M, Arce-Rodríguez A, De Las Heras A, Páez-Espino AD, Durante-Rodríguez G, Kim J, Nikel PI, Platero R, De Lorenzo V. 2013. The Standard European Vector Architecture (SEVA): A coherent platform for the analysis and deployment of complex prokaryotic phenotypes. Nucleic Acids Res 41:666–675.

60. Bagdasarian M, Lurz R, Rückert B, Franklin FCH, Bagdasarian MM, Frey J, Timmis KN. 1981. Specific-purpose plasmid cloning vectors II. Broad host range, high copy number, RSF 1010-derived vectors, and a host-vector system for gene cloning in Pseudomonas. Gene 16:237– 247.

61. Martínez-García E, Nikel PI, Aparicio T, De Lorenzo V. 2014. Pseudomonas 2.0: genetic upgrading of P. putida KT2440 as an enhanced host for heterologous gene expression. Microb Cell Fact 13:1–15.

62. Stover CK, Pham XQ, Erwin AL, Mizoguchi SD, Warrener P, Hickey MJ, Brinkman FSL, Hufnagle WO, Kowallk DJ, Lagrou M, Garber RL, Goltry L, Tolentino E, Westbrock-Wadman S, Yuan Y, Brody LL, Coulter SN, Folger KR, Kas A, Larbig K, Lim R, Smith K, Spencer D, Wong GKS, Wu Z, Paulsen IT, Relzer J, Saler MH, Hancock REW, Lory S, Olson M V. 2000. Complete genome sequence of Pseudomonas aeruginosa PAO1, an opportunistic pathogen. Nature 406:959– 964.

