## Supplementary material for "A SEVA-based, CRISPR-Cas3-assisted genome engineering approach for *Pseudomonas* with efficient vector curing": Figure S

### Supporting information

#### Supplementary Tables

Table S1: All primers used in this work. Red: overhang for ligation in pCas3cRh with BsaI; yellow: BsaI recognition site; blue: SapI recognition site; grey: BsaI/SapI restriction site; green: U nucleotide for USER cloning

| **Name** | **Sequence** |
| --- | --- |
| *Vector construction – pSEVA52-oriT* | |
| oriT_spacer_F | GAAACGCACGATATACAGGATTTTGCCAAAGGGTTCGTGG |
| oriT_spacer_R | GCGACCACGAACCCTTTGGCAAAATCCTGTATATCGTGCG |
| oriTcas3_SEVA_F | GACGGTCTCCCTCTACTAGTCTTGGACTCCTG |
| oriTcas3_SEVA_R | GTCGGTCTCCGGTGTTAATTAAAGGCATCAAATAAAACG |
| oriT_cas3_F | GACGGTCTCCCACCACAATTCAGCAAATTGTGAAC |
| oriT_cas3_R | GTCGGTCTCCAGAGCACTAGGTTTCAATCCAC |
| *Vector assembly – pCas3cRh with spacers* | |
| pCas3cRh_F | CAGGAAATGCGGTGAGC |
| pCas3cRh_R | GAGCAGCTAATTCACCGC |
| PP5388_spacer_F | GAAACAGATCATGGTAACCCCGGCCGCTGGAGCCATTTCG |
| PP5388_spacer_R | GCGACGAAATGGCTCCAGCGGCCGGGGTTACCATGATCTG |
| FleS_spacer_F | GAAACAACCAGATGTCCAGCCAGCTCAGCGAGTCCTACAG |
| FleS_spacer_R | GCGACTGTAGGACTCGCTGAGCTGGCTGGACATCTGGTTG |
| PrpL_spacer_F | GAAACTACAACACCACCCAGTGCTACGGCGACGCCTCGAG |
| PrpL_spacer_R | GCGACTCGAGGCGTCGCCGTAGCACTGGGTGGTGTTGTAG |
| PA2560_spacer_F | GAAACGCCTGCTGGACGAGGTGCAATACCCGGATGCGCCG |
| PA2560_spacer_R | GCGACGGCGCATCCGGGTATTGCACCTCGTCCAGCAGGCG |
| *Vector assembly – HDR templates in pSEVAX31* | |
| SEVA_PS1 | AGGGCGGCGGATTTGTCC |
| SEVA_PS2 | GCGGCAACCGAGCGTTC |
| pSNW_seq_F | TGTAAAACGACGGCCAGT |
| pSNW_seq_R | CTTTACACTTTATGCTTCCGG |
| pSEVA231 _F | GACGGTCTCCAGTCACTAGTCTTGGACTCCTG |
| pSEVA231 _R | GTCGGTCTCCCTAGTTAATTAAAGGCATCAAATAAAACG |
| pSEVA131_FleS_F | TACGGTCTCAAGTCACTAGTCTTGGACTCCTG |
| pSEVA131_Fles_R | GGGGGTCTCCGGACTTAATTAAAGGCATCAAATAAAACG |
| pSEVA131_PrpL_F | GACGGTCTCCGGGCACTAGTCTTGGACTCCTG |
| pSEVA131_PrpL_R | GTCGGTCTCCTTACTTAATTAAAGGCATCAAATAAAACG |
| pSEVA131_PA2560_F | GACGGTCTCCTGGCACTAGTCTTGGACTCCTG |
| pSEVA131_PA2560_R | GTCGGTCTCCGAGCTTAATTAAAGGCATCAAATAAAACG |
| pSNW2_F | ATAGGTCTCAAGTCGACCTGCAGGCATGCAAGC |
| pSNW2_R | ATAGGTCTCACTAGAGGATCCCCGGGTACCGAGC |
| PP5388_HAup_F | ATAGGTCTCACTAGTGACCGCACCGTTGCTCTCTCTCTTCG |
| PP5388_HAup_R | ATAGGTCTCATAGATGAATCGCCCTGACTTC |
| PP5388_HAdn_F | ATTGGTCTCAGGCTGATCATGGTAACCCCGGCCGC |
| PP5388_HAdn_R | ATAGGTCTCAGACTGCCCTGGTCGACGATATTG |
| Trrnb_F | ATAGGTCTCATCTAATTTGTCCTACTCAGGAGAGCG |
| T0term_R | CTTGGTCTCTAGCCCTGGATTCTCACCAATAAAAAAC |
| FleS_HAup_F | TTAGGTCTCCGTCCGCGAACTGGCGAACC |
| FleS_HAup_R | TATGGTCTCCTTGCGTTTCTCTCGGTCGGCGCGACG |
| FleS_HAdn_F | ATAGGTCTCCAACGCACCCCATGGCAGCCAAAGTCC |
| FleS_HAdn_R | AACGGTCTCTGACTTCCTTGCCGGTCCCGGACTC |
| PrpL_HAup_F | GACGGTCTCCGTAATGGTAAGCGGCCAG |
| PrpL_HAup_R | GTCGGTCTCCCCGAATCGACTCCTTCAG |
| PrpL_HAdn_F | GACGGTCTCCTCGGCCAGCCAGGCTATCG |
| PrpL_HAdn_R | GTCGGTCTCCGCCCGCGAGCATTCCTCCTG |
| PA2560_HAup_F | GACGGTCTCCGCTCTCCAGTATTTCGTC |
| PA2560_HAup_R | GTCGGTCTCCGCGAGTCAGATCCTTCG |
| PA2560_HAdn_F | GACGGTCTCCTCGCCGACGCCAGGAGCAC |
| PA2560_HAdn_R | GTCGGTCTCCGCCAAGGCGCAAGCCTG |
| *Verification of genomic modifications* | |
| PP5388_up | ATTCTTTGGAGCAACGGAAC |
| PP5388_dn | CAGCTGCAAGTGGAGATC |
| FleS_up | AGAAGATGATCGAGGAC |
| FleS_dn | GAACCTTTCTCGTGGC |
| PrpL_up | CCACGTCAGCGGCAAG |
| PrpL_dn | GGCTCTGCCGTTTCAC |
| PA2560_up | GTTGCCTGCCTTGTC |
| PA2560_dn | GACGCCTTGCTTCTC |
| *Vector construction – pCas3-XX and pSEVAX3-GG vector sets* | |
| pCas3_Ab_F | AACAGGAGUAGCGTCAGACCCCGTAGAAAAG |
| pCas3_Ab_R | ATCCATCCUGCGTGAGCGCATACGCTACTTG |
| Ab_F | ACTCCTGTUGATAGATCCAGTAATGACCT |
| Ab_R | AGGATGGAUATACCGAAAAAATCGCTATAA |
| pSX31_GG_F | AGGTCTCTGAAUCCCCGGGTACCGAGC |
| pSX31_GG_R | AGTGGTCTCGUCGCCCGGCAAAACCGGGCGTGGATTGAAACGCTTGCGGCCGCGTCG |
| P14g-BCD2-GFP_F | ATTCAGAGACCUGCCCATTGACAAGGC |
| P14g-BCD2-GFP_R | ACGAGACCACUAGTCATTTATTTGTAGAG |
| pSX31_Ab_F/R | ATCCATCCUTTTTCGCACGATATACAGG |
| pSX31_Ab_F/R | AACAGGAGUCCAAGACTAGTCGC |
| ApR_BsaI_F | AGCCAAUCGACTGGCGAGCGGCATC |
| ApR_BsaI_R | ATTGGCUAAGACCGCGGTCCGCGCGT |

Table S2: All vectors used in this work.

| **Name** | **Relevant features** | **Reference** |
| --- | --- | --- |
| pCas3cRh | *oriT*; *oriV(pRO1600/ColE1)*; RhaRS/*PrhaBAD::gRNA-cas3-cas5-cas8-cas7*; MCS; Gm^R^ | (58) |
| pCas3cRh-PP_5388 | pCas3cRh derivative with a PP5388-targeting spacer | This work |
| pCas3cRh-FleS | pCas3cRh derivative with a FleS-targeting spacer | This work |
| pCas3cRh-PrpL | pCas3cRh derivative with a PrpL-targeting spacer | This work |
| pCas3cRh-PA2560 | pCas3cRh derivative with a PA2560-targeting spacer | This work |
| pCas3cRh-oriT | pCas3cRh derivative with an oriT-targeting spacer | This work |
| pCas3-Amp | *oriT*; *oriV(pRO1600/ColE1)*; RhaRS/*PrhaBAD::gRNA-cas3-cas5-cas8-cas7*; MCS; Amp^R^ | This work |
| pCas3-Km | *oriT*; *oriV(pRO1600/ColE1)*; RhaRS/*PrhaBAD::gRNA-cas3-cas5-cas8-cas7*; MCS; Km^R^ | This work |
| pCas3-Sm | *oriT*; *oriV(pRO1600/ColE1)*; RhaRS/*PrhaBAD::gRNA-cas3-cas5-cas8-cas7*; MCS; Sm^R^ | This work |
| pCas3-Gm | *oriT*; *oriV(pRO1600/ColE1)*; RhaRS/*PrhaBAD::gRNA-cas3-cas5-cas8-cas7*; MCS; Gm^R^ | This work |
| pCas3-Apr | *oriT*; *oriV(pRO1600/ColE1)*; RhaRS/*PrhaBAD::gRNA-cas3-cas5-cas8-cas7*; MCS; Apr^R^ | This work |
| pSEVA231 | SEVA vector, *oriT*; *oriV(pBBR1)*; MCS; Km^R^ | (59) |
| pSEVA23-PP_5388 | pSEVA231 derivative with PP5388(HA1)-T1-*p14c-BCD22-phi15lys(G3RQ)-*T0-PP0013(HA2) | This work |
| pSNW2 | Integration vector; *oriT*; *oriV(R6K)*; *P_14g_-BCD2-msfGFP* selection marker; MCS*,* Km^R^ | (36) |
| pSNW2-PP_5388 | pSNW2 with PP5388(HA1)-T1-*p14c-BCD22-phi15lys(G3RQ)-*T0-PP0013(HA2) | This work |
| pSEVA131 | SEVA vector, *oriT*; *oriV(pBBR1)*; MCS; Amp^R^/Cb^R^ | (59) |
| pSEVA13-FleS | pSEVA131 derivative with FleS(HA1)-FleS(HA2) | This work |
| pSEVA13-PrpL | pSEVA131 derivative with PrpL(HA1)-PrpL(HA2) | This work |
| pSEVA13-PA2560 | pSEVA131 derivative with PA2560(HA1)-PA2560(HA2) | This work |
| pSEVA521 | SEVA vector, *oriT*; *oriV(RK2)*; MCS; Tc^R^ | (59) |
| pSEVA52-oriT | pSEVA521 derivative with *PrhaBAD::CRISPRrepeat-oriTspacer-CRISPRrepeat* | This work |
| pSEVA13-GG | SEVA vector, *oriT*; *oriV(pBBR1)*; GoldenGate compatible with BsaI sites flanking a *P_14g_-BCD2-msfgfp* reporter cassette; Amp^R^/Cb^R^ | This work |
| pSEVA23-GG | SEVA vector, *oriT*; *oriV(pBBR1)*; GoldenGate compatible with BsaI sites flanking a *P_14g_-BCD2-msfgfp* reporter cassette; Sm^R^/Sp^R^ | This work |
| pSEVA43-GG | SEVA vector, *oriT*; *oriV(pBBR1)*; GoldenGate compatible with BsaI sites flanking a *P_14g_-BCD2-msfgfp* reporter cassette; Km^R^ | This work |
| pSEVA63-GG | SEVA vector, *oriT*; *oriV(pBBR1)*; GoldenGate compatible with BsaI sites flanking a *P_14g_-BCD2-msfgfp* reporter cassette; Gm^R^ | This work |
| pSEVA83-GG | SEVA vector, *oriT*; *oriV(pBBR1)*; GoldenGate compatible with BsaI sites flanking a *P_14g_-BCD2-msfgfp* reporter cassette; Apr^R^ | This work |
| pBG42 | pBG derivative with P14g-BCD2-msfGFP | (48) |

Table S3: All strains used in this work.

| **Name** | **Description** | **Reference** |
| --- | --- | --- |
| *E. coli* TOP10 | Intermediate host for vector cloning (Invitrogen^TM^); F- *mcrA* Δ( *mrr-hsd*RMS-*mcr*BC) Φ80*lac*ZΔM15 Δ *lac*X74 *rec*A1 *ara*D139 Δ( *araleu*)7697 *gal*U *gal*K *rps*L (StrR) *end*A1 *nup*G | - |
| *P. putida* KT2440 | Derivative of *P. putida* mt-2 lacking the TOL plasmid | (60) |
| *P. putida* SEM11 | Genome reduced derivative of *P. putida* KT2440 with deletion of prophages, ꞵ-lactamases, *benABCD* and *pvdD* | (37, 61) |
| *P. putida* KT-phi15lys | Derivative of *P. putida* KT2440 with a T1-p14c-BCD22-phi15lys(G3RQ)-T0 integration in the PP5388 locus | This work |
| *P. putida* S-phi15lys | Derivative of *P. putida* SEM11 with a T1-p14c-BCD22-phi15lys(G3RQ)-T0 integration in the PP5388 locus | This work |
| *P. aeruginosa* PAO1 |  | (62) |
| *P. aeruginosa* PAO1 Δ*fleS* | Derivative of *P. aeruginosa* PAO1 with a deletion of the *fleS* gene | This work |
| *P. aeruginosa* PAO1 Δ*prpL* | Derivative of *P. aeruginosa* PAO1 with a deletion of the *prpL* gene | This work |
| *P. aeruginosa* PAO1 *Δpa2560* | Derivative of *P. aeruginosa* PAO1 with a deletion of the *pa2560* gene | This work |

Table S4: All sequences of homology arms and inserts used in this work. HA: homology arm, up: upstream, dn: downstream

| **Name** | **Sequence** |
| --- | --- |
| PP_5388 HA up | TGACCGCACCGTTGCTCTCTCTCTTCGTGATTCCCGCAGCGTATTGGCTGGTCCGACGCCGCGATCTTGTAGTACCTCATAATTCCACACCAGGAGACATCCGATGAAAAAGCTCTACCTCAGCATTGCACTGCTCTTTGCCTTCGCATCAGGCGCGCAAGCCCAAGACTCCATGGCCGGGATGAGCATGGATGGAATGGATATGAAGGAAACCCAATCAGCACCTTCCGCTCATGCAGAAGGGACGGTAAAGGCAATTGACGCCCAAGGCGGCACAGTGACCCTGATGCATGGACCGGTTGCTGCGTTGAAATGGCCGGCCATGACCATGGCCTTCAAAGCCTCTGCGCAACAGCTCGATGGATTGAAGGTGGGAGACAACGTAGAGTTTGATTTCCGGATGGATGGCAGCACGGCAACGATTGTTGATATTCGCAAACAGTAATTCGCCTAATTATCCTGTCTGCCAACGCTTGATTCGCCCATGACCTCCTGACGCCTCCTAGCAACTGGAAAAGCTTGGAAGCATCTAGGAAGTCAGGGCGATTCA |
| PP_5388 HA dn | GATCATGGTAACCCCGGCCGCTGGAGCCATTTCTGAAGTACTGACACGGCGCTGAAGCGATGTGGCAGTTCCGCCTAAGGCGGCGAGCAGCAGCGAGAAAAGATCGTCACCCCAAGTTTTCTTCTTGCAGGCGACTGAATAGAGGAGAAATCGAGATAACAGCTCGCAAGGCATCCTGATTGGCGCCCGACCATGCCTGATCCTGTACATCATTGAGCACGTGCTGTGTCGGAATTGGCCGAAACGCTTGCGTCGCGGGGTCGTAGACGTAAACGTGAAGACCTATTTCAATCGCTCGCGAGGCCTGACGGTGCCAGTTAAAGTTGCCACCATTCGTCATCGTGTCCGCAGCTACGAGGATGTCGTAGTCATTAACGATGTAGCCGAAAAGTACATGCTGCGAGATATCGCGGAGAGCTAGAGAATGACGCCTGCTCGCCGATCGCCAGACTAGGTTGCGTGCAAAGTGTCGCCTACTGACCCTTGGGTCATCCAAAAGCGTGACCTTGTCGTAATAGGCTACTGACAAATCAATATCGTCGACCAGGGC |
| *P14c-BCD22-phi15lys(G3RQ)* cassette | ATTTGTCCTACTCAGGAGAGCGTTCACCGACAAACAACAGATAAAACGAAAGGCCCAGTCTTTCGACTGAGCCTTTCGTTTTATTTGATGCCTTTAATTAATCTAGTGAATTGACATGTCAATTTTTATGTTGTATAATATAACTAGCAGGCCCAAGTTCACTTAAAAAGGAGATCAACAATGAAAGCAATTTTCGTACTGAAACATCTTAATCATGCCTAGGAAGTTTTCTAATGGCTCGTCAAGTCAAATTTAAGGAGCGCTTGAGTACCAAGATGATCGTTGTGCACTGTTCGGCCACCAAGGCCAGCATGGACATCGGTCGTAAAGAGATCCAAATGTGGCACGTTCAGCAAGGCTGGCTGGCCATTGGGTACCACCTCGTGATTCGTCGCGACGGTACCATCGAGCAAGGCCGACCACACAAGGCCATCGGGTCCCACGTTAAGGGTCACAACAGTGACTCCATCGGGATCTGCCTCGTGGGCGGTATCGACGACTCGGGGAAACCTGAGGACAACTTCACGGACCAGCAAAAGGCTGCGCTGAGCGGCCTGCTGTGGGACATGACGCAATCTGGCGTGACCTATGGGGACACCTATAAGGAGCTGCCAGTGGTTGGTCACCGTGATCTCGATAGTGGTAAAGCCTGCCCGAGCTTCGATGTGAAGGCATGGTGGGCCGCTCAGATCAATTAACACTAGTCTTGGACTCCTGTTGATAGATCCAGTAATGACCTCAGAACTCCATCTGGATTTGTTCAGAACGCTCGGTTGCCGCCGGGCGTTTTTTATTGGTGAGAAT |
| FleS HA up | CCGCGAACTGGCGAACCTGGTGGAGCGCCTGGCGATCATGCATCCCTACGGGGTGATCGGGGTCGGCGAACTGCCGAAGAAATTCCGCCATGTCGACGACGAGGACGAGCAACTCGCCAGCAGCCTGCGCGAAGAGCTGGAAGAGCGCGCGGCGATCAACGCCGGGCTGCCGGGAATGGACGCGCCGGCGATGCTGCCGGCCGAAGGCCTGGACCTCAAGGACTACCTGGCCAACCTCGAGCAGGGCCTGATCCAGCAGGCCCTCGACGACGCCGGCGGAGTGGTCGCGCGGGCCGCCGAACGCCTGCGCATCCGCCGCACCACGCTGGTAGAGAAGATGCGCAAGTACGGCATGAGCCGGCGTGACGACGACCTGTCGGATGATTGACAGGTCGTTTCGCAACGCTTTGATTTTCAAATGAAAAAAATTTAGGCACGGGTATTGCTATATCTCCGTCGACCGACAGAACCATGACGTCGCGCCGACCGAGAGAAACG |
| FleS HA dn | ACCCCATGGCAGCCAAAGTCCTGCTGGTCGAAGACGACCGCGCACTACGCGAAGCCCTCAGCGACACCCTGCTGCTGGGCGGTCACGAGTTCGTCGCCGTGGACTCGGCGGAGGCGGCGCTGCCGGTCCTGGCCCGCGAAGCCTTCAGCCTGGTGATCAGCGACGTGAACATGCCGGGCATGGACGGACACCAGTTGCTCGGCCTGATCCGTACACGCTACCCGCACCTGCCGGTGTTGCTGATGACCGCCTACGGCGCGGTCGATCGCGCCGTCGAGGCGATGCGCCAGGGCGCCGCCGACTACCTGGTCAAGCCGTTCGAGGCGCGGGCGCTGCTCGACCTGGTGGCGCGCCATGCGCTGGGCCAGTTGCCGGGCAGCGAGGAGGATGGTCCGGTGGCCCTGGAGCCGGCCAGCCGGCAGTTGCTGGAACTGGCCGCGCGGGTCGCGCGCAGCGATTCCACCGTGCTGATCTCCGGCGAGTCCGGGACCGGCAAGGAAG |
| PrpL HA up | GTAATGGTAAGCGGCCAGGGCTCTATATATGCCCCGCGATTATAGAGATGACTTATACGACCATAAGCCGGATATCCTGGAATTCAACCAAGCCAACATCGGAACGGTTTCCGAGCGGTGAATGGAAGATTCACCCAGCCTTATATTCGGCAGTGGTCAAACCGGATAACTTGAAAATCAGGATTTCGTTAACCCGATCACATCGACAGCTGCACCAGTAAAGCACTCATGCACTACATCCTGCACCTCTCTGAAGCAAACCGAAGGCTCTGCAGAGCCACTCCAGACCAAACTTTAGTTGGTAGAGAGAGCAATCCAACATCAATGGCAAGTGAAGGAAATAGCTATCTATTCTGCTATCCACAAACGCTTCTTCATATTTAATAGCTAGCAGAAACGAATATAAAAAATATTTGAACGACTCCGTCACACCTGCATATCTTTCGCCACGCGAACGATTGAGGAAGTTGCCCTCCCAAAAAACTGAAGGAGTCGATTC |
| PrpL HA dn | GGCCAGCCAGGCTATCGATCGCGCACCGGCGGATAACCGCGCAGCGGTTATTCGCCCTACGCCCGGATTGGCGCCTGGAAGTGCGGCTTCTTTCGTCGTCCGGGACAGGTAGGGCGCATAACGCCAACGGCGTTATCCGCCGTCTATCCCGCGCAATGGTTATTCGCCCCACGCCCGGATTGGCACCTGGAAGTGCGGCTTCTTTCGTCGTCCGGGACAGGTAGGGCGCATAACGCCAACGGTGTTATCCGCCGTTTATCCCGCGCAATGGTTATTCGCCCCACGCCTGGATCGGCGCGTGGAAGAGGGGCGTTATGCGCCGCCGTACGTTCAGTCCTGGCGGCTGGTGACTTCCAGCAGGTGGTAGCCGAACTGGGTCTTCACCGGCCCCTGGACCACGTTCAGCGGCGCGCTGAAGACCACCTGGTCGAACTCGCGGACCATCTGGCCGGGGCCGAACGAGCCCAGGTTGCCGCCGTCGCGGCCGGAGGGGCAGGAGGAATGCTCGCGGGC |
| PA_2560 up | GAGTCAGATCCTTCGACTGGCACAATGTGACATTCGTCACAGAGATTAACCGAAGCCCCGCAGAACATCCATGCCAGCTATTTCCCTTCGTCGGCTCTGGCCCGGCCCCTGCGTCGTCCTCAAGTTGTCAGATCCGCTGTCGATACTCTGCGAGCCGGAACACGCGCCATGCCCGCCTCGCCGGGCCACAGGGACGTCCTCGGCTGCCTCGTCGCAGCCTGCGTGCCGGTCCAACCTGGCAATCCATCGAGGCGTTCCATGCTGCAACAATCCCTACGTGCGCAAATCCTTGTCCTGCTCGGCGGCAGCCTGGCGGCGCTGCTACTCATAGCCCTGGCCTGCTTCGGCTCGCTGACCGGCGACGTACGCGCCTACCGCGAGCTGCTCGGCGGCCCCGTGCGGGCGGCGCAACTGATCGACGAGGCCAACCTGCAATTCCGCGGCCAGGTCCAGGAATGGAAGAACGTCCTGCTGCGCGGACGCCAGACGGAGGCCCAGACGAAATACTGGA |
| PA_2560 dn | GCCAAGGCGCAAGCCTGAACAAGCTAGCCGACTTGGCGAGGAACGCACAAATTTCATCGGCTGGCGTACAGGTATGGGACATCGGTACCAATAGCGAAGTCGCGCAATCCGTCAGTTTTTTTTGAATTTTTGCCATTGGAAAAAGTCTGTTCAGCGCAGCGTTTTTCCAAGTCCCATCCGGCCCGCAAGGAGAACCCATGGCTGAACCCCAGGACAAGTACACCCGGCGCACAGGCAGGACCTGGGCGGACGACCAGGCGACCTACAACCGTCTGCGCGAAGAAGCCGACGCCGCTCGCCAGAAGCTGCGCGAAAGCGGCTACAGCGGCGCCGAGTACGACCAGTTGCGTCAAGCCGCCTTCGATCTCAACCGCAAGGCCAACCAGTACTGGGAGCAGATGCTCAGCGACCTGCGCCAGGAAGACTGATCAGCGCCACGCCGGGAGACTGCCGCTCTCGGCCCGCACACCCCCTCCACTCTTTGCGTGCTCCTGGCGTCGGC |

Table S5: SNP analysis of P. putida KT-phi15lys, replicate 1.

| **Reference** | **Nucleotide position** | **Type** | **Reference** | **Alteration** | **Evidence** |
| --- | --- | --- | --- | --- | --- |
| genome | 533982 | snp | A | G | G:31 A:0 |
| genome | 1126645 | ins | A | AC | AC:21 A:0 |
| genome | 1720346 | snp | C | T | T:27 C:0 |
| genome | 4586056 | ins | A | AC | AC:19 A:0 |

Table S6: SNP analysis of P. putida KT-phi15lys, replicate 2.

| **Reference** | **Nucleotide position** | **Type** | **Reference** | **Alteration** | **Evidence** |
| --- | --- | --- | --- | --- | --- |
| genome | 533982 | snp | A | G | G:52 A:0 |
| genome | 1126645 | ins | A | AC | AC:20 A:0 |
| genome | 1720346 | snp | C | T | T:37 C:0 |
| genome | 4586030 | complex | CTGC | TCGCG | TCGCG:11 CTGC:0 |

Table S7: SNP analysis of P. putida S-phi15lys, replicate 1.

| **Reference** | **Nucleotide position** | **Type** | **Reference** | **Alteration** | **Evidence** |
| --- | --- | --- | --- | --- | --- |
| genome | 1146407 | snp | A | C | C:10 A:0 |
| genome | 5200648 | snp | T | C | C:14 T:0 |

Table S8: SNP analysis of P. putida S-phi15lys, replicate 2.

| **Reference** | **Nucleotide position** | **Type** | **Reference** | **Alteration** | **Evidence** |
| --- | --- | --- | --- | --- | --- |
| genome | 1146407 | snp | A | C | C:12 A:0 |
| genome | 5200648 | snp | T | C | C:17 T:0 |

#### Supplementary Figures

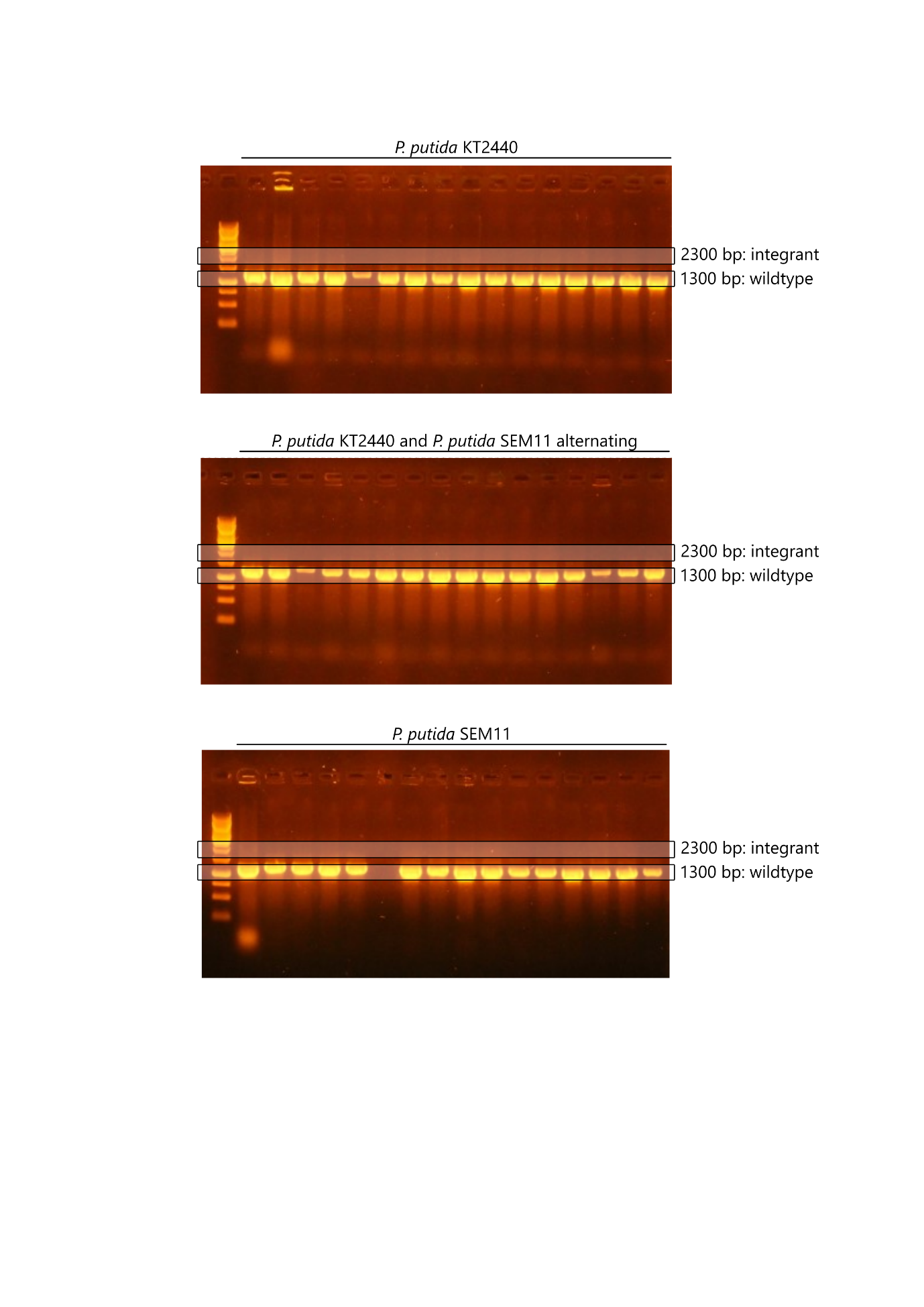

Figure S1: Integration of P_14c_-BCD22-phi15lys(G3RQ) in the PP_5388 locus in P. putida KT2440 and P. putida SEM11 with Cas3-based engineering. A PCR screen was performed on co-transformants (no induction) with genome-binding primers located outside of the homology arms. The first lane on each gel contains the commercial 1 kb GeneRuler (Thermo Scientific). Expected amplicon length of wildtype colonies and integrants are indicated on each gel.

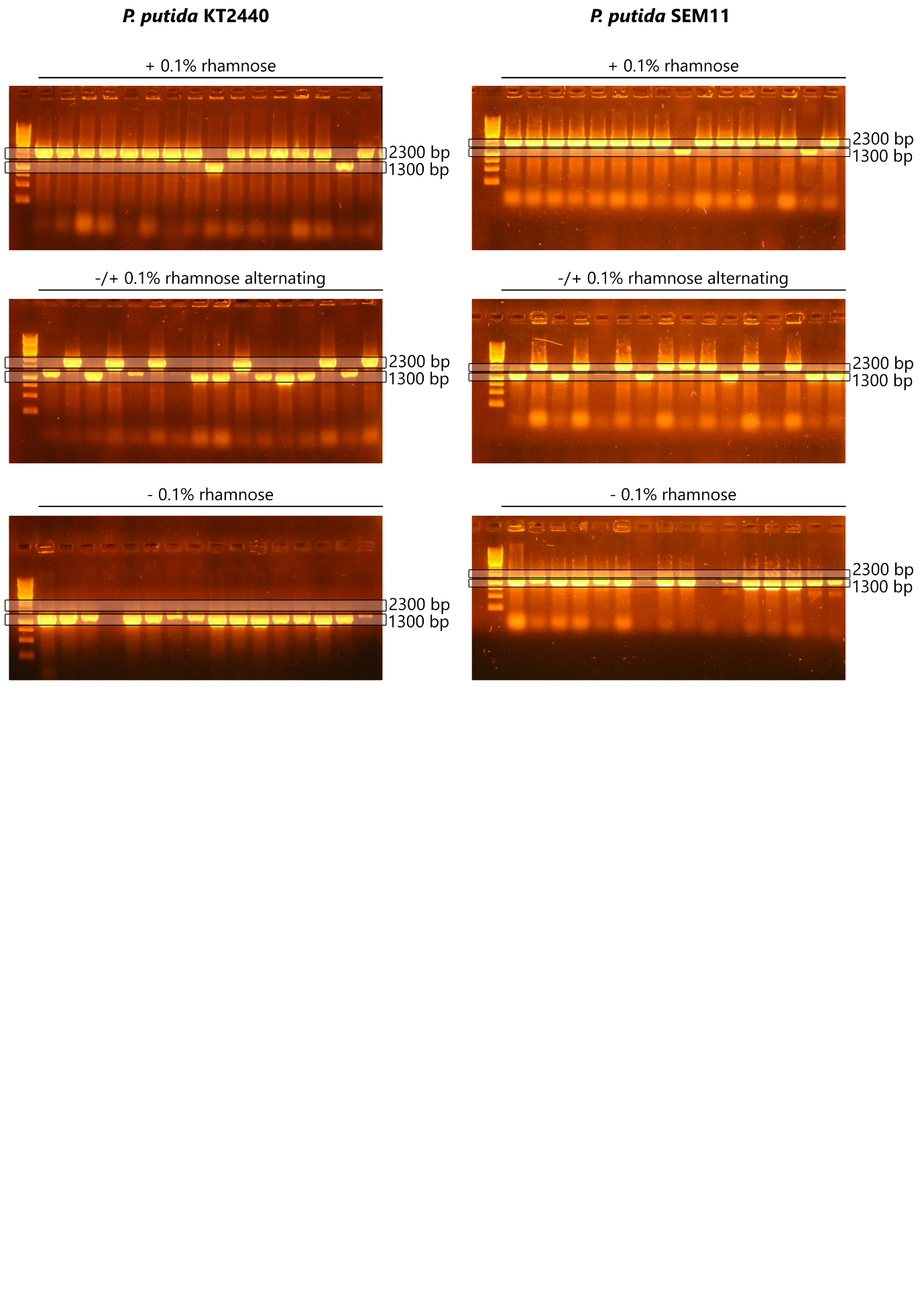

Figure S2: Integration of P_14c_-BCD22-phi15lys(G3RQ) in the PP_5388 locus in P. putida KT2440 (left) and P. putida SEM11 (right) with Cas3-based engineering. A PCR screen was performed on co-transformants (with and without overnight induction with 0.1% rhamnose) with genome-binding primers located outside of the homology arms. The first lane on each gel contains the commercial 1 kb GeneRuler (Thermo Scientific). Expected amplicon length of wildtype colonies (1300 bp) and integrants (2300 bp) are indicated on each gel.

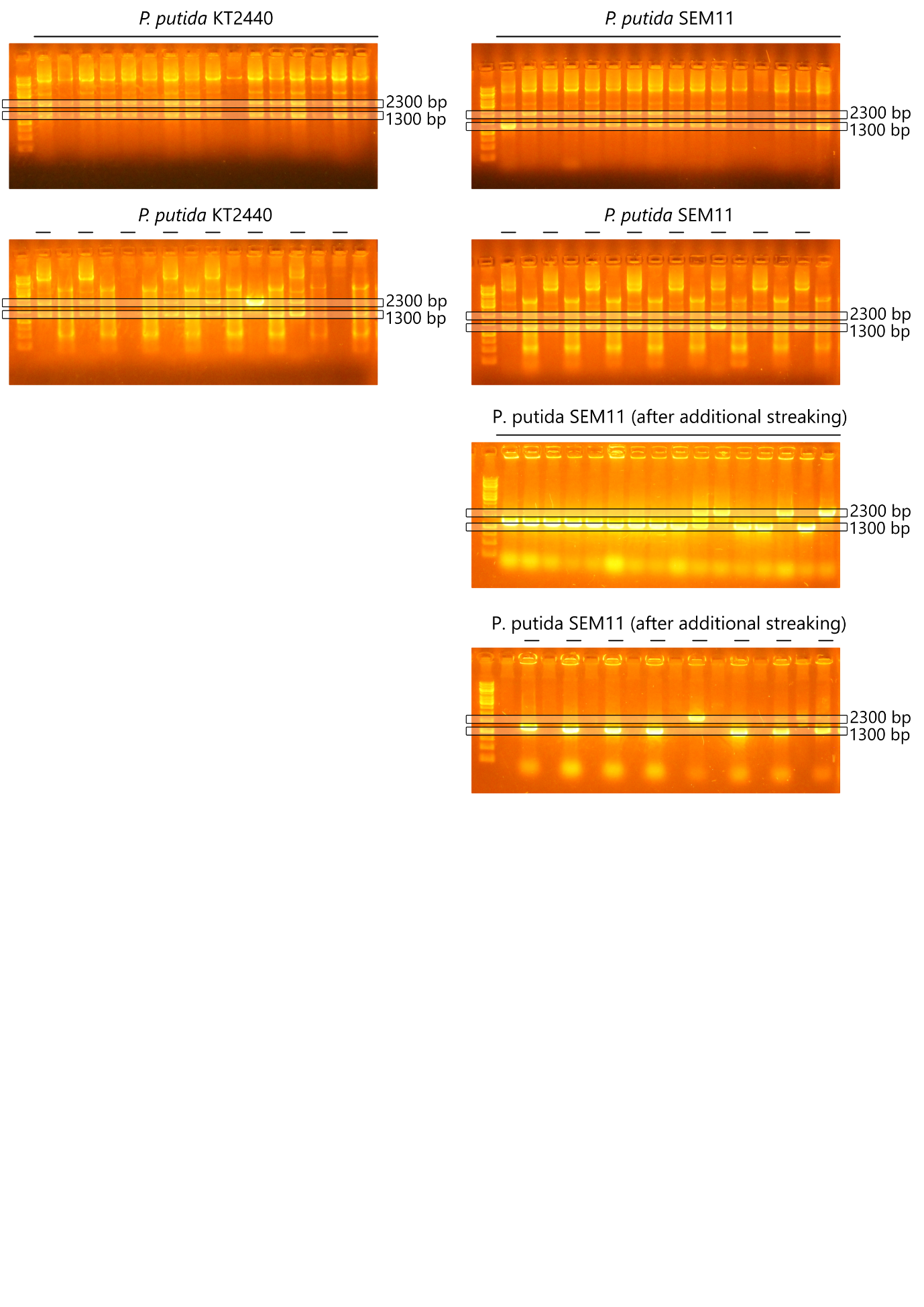

Figure S3: Integration of P_14c_-BCD22-phi15lys(G3RQ) in the PP_5388 locus in P. putida KT2440 (left) and P. putida SEM11 (right) with homologous recombination. A PCR screen was performed on co-transformants with genome-binding primers located outside of the homology arms. The first lane on each gel contains the commercial 1 kb GeneRuler (Thermo Scientific). Expected amplicon length of wildtype colonies (1300 bp) and integrants (2300 bp) are indicated on each gel.

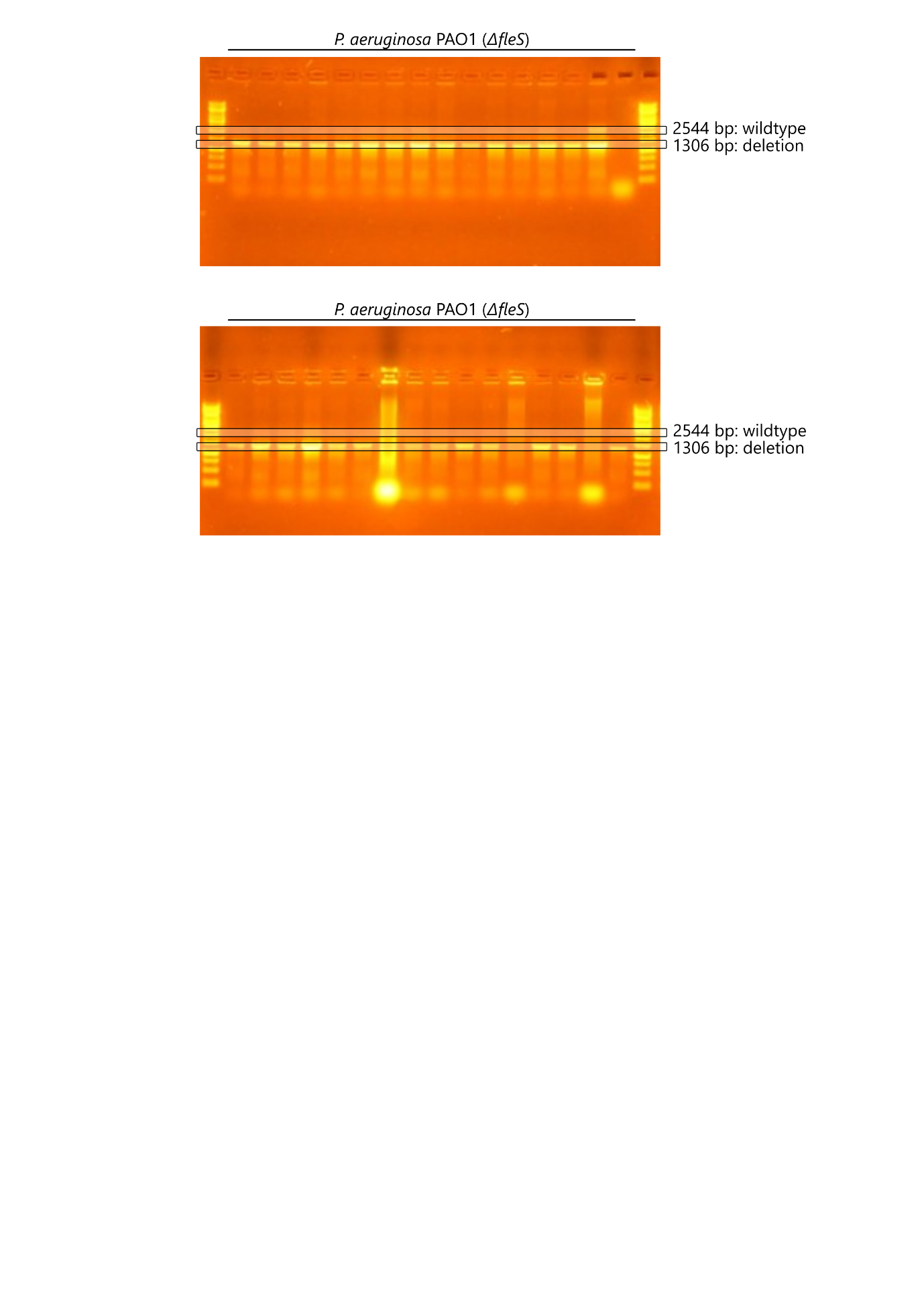

Figure S4: Deletion of fleS in P. aeruginosa PAO1 with Cas3-based engineering. A PCR screen was performed on co-transformants with genome-binding primers located outside of the homology arms. The first lane on each gel contains the commercial 1 kb GeneRuler (Thermo Scientific). Expected amplicon length of wildtype colonies (2544 bp) and deletion mutants (1306 bp) are indicated on each gel.

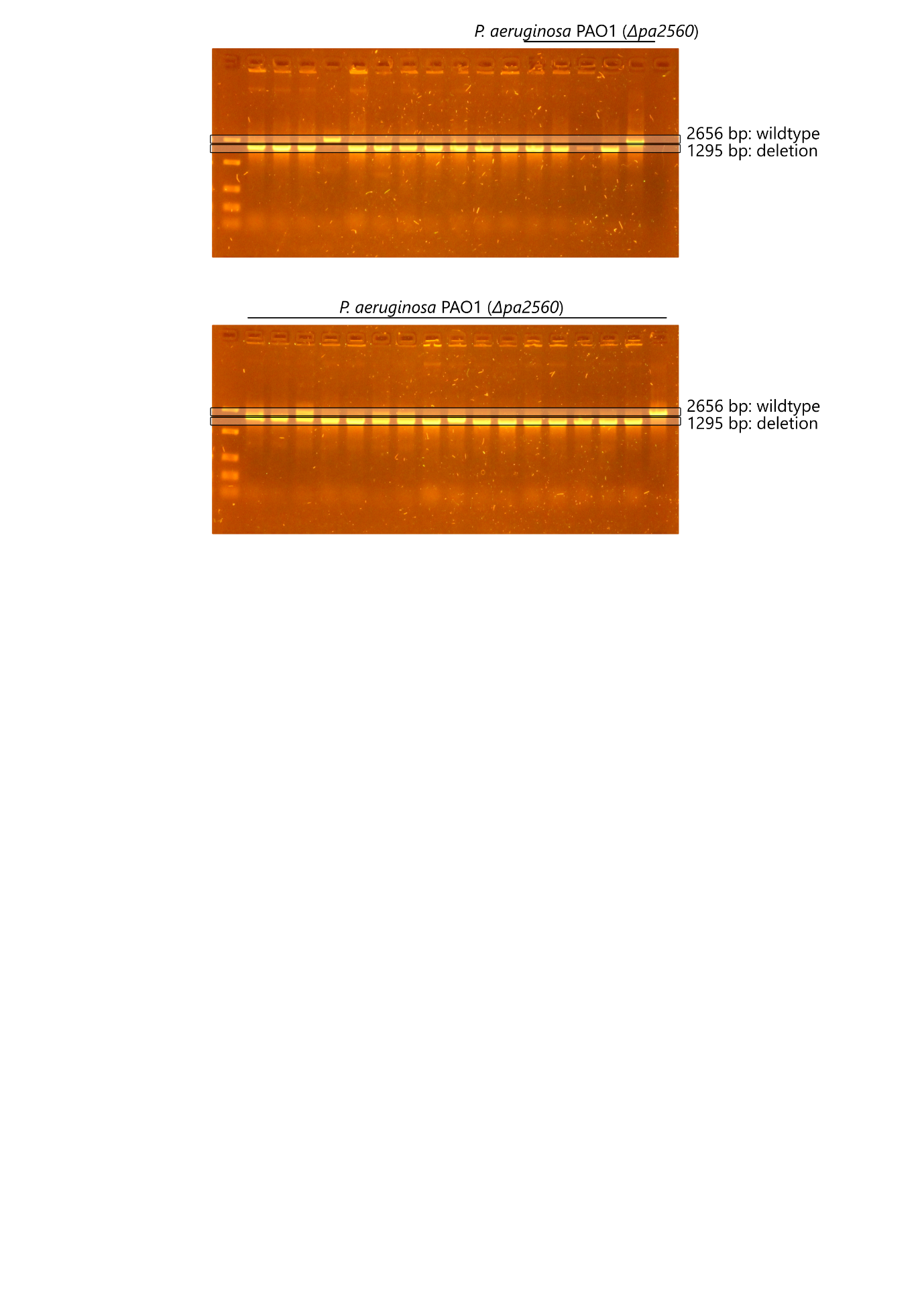

Figure S5: Deletion of PA_2560 in P. aeruginosa PAO1 with Cas3-based engineering. A PCR screen was performed on co-transformants with genome-binding primers located outside of the homology arms. The first lane on each gel contains the commercial FastRuler Low Range DNA ladder (Thermo Scientific). Expected amplicon length of wildtype colonies (2656 bp) and deletion mutants (1295 bp) are indicated on each gel.

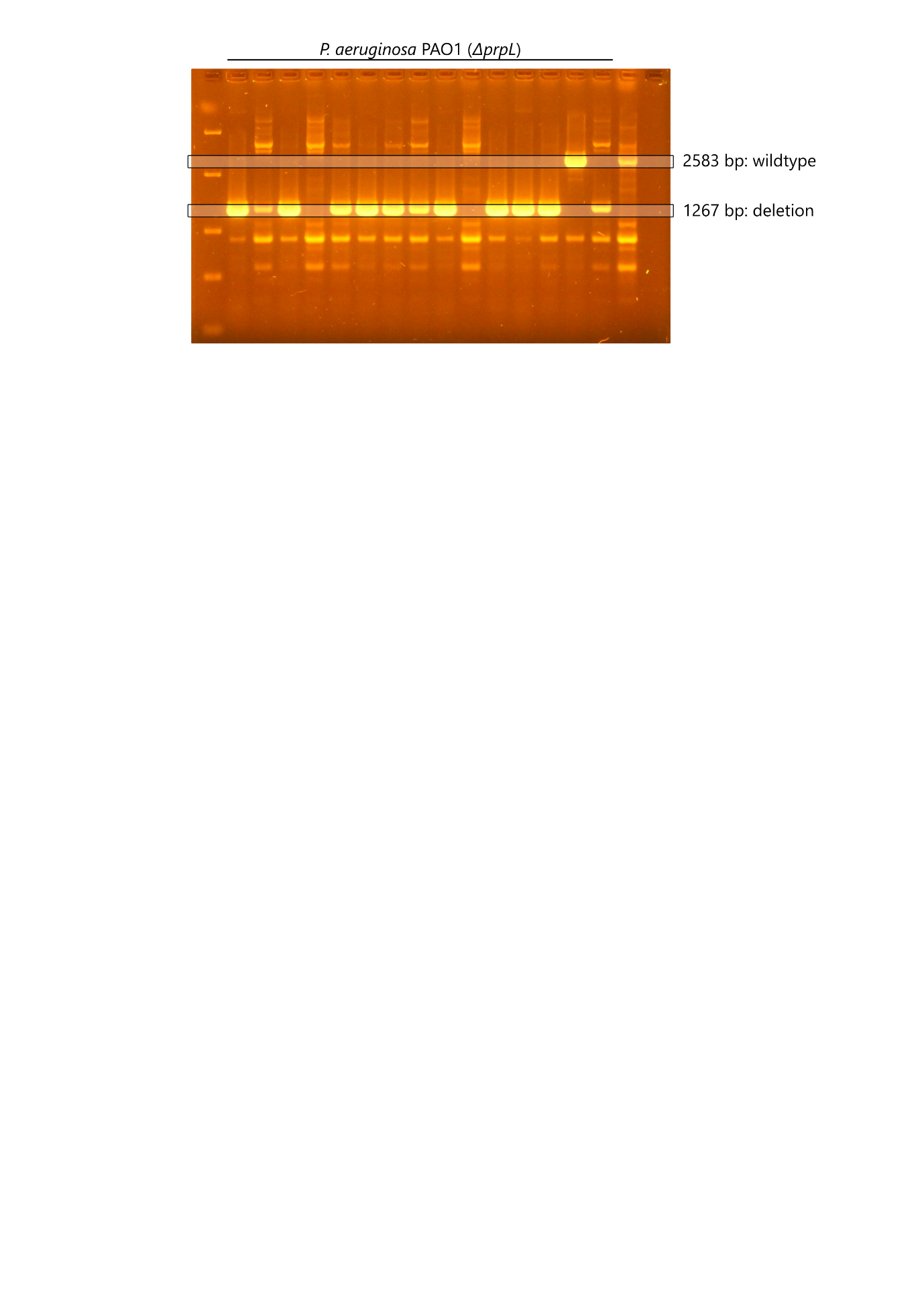

Figure S6: Deletion of prpL in P. aeruginosa PAO1 with Cas3-based engineering. A PCR screen was performed on co-transformants with genome-binding primers located outside of the homology arms. The first lane on each gel contains the commercial FastRuler Low Range DNA ladder (Thermo Scientific). Expected amplicon length of wildtype colonies (2583 bp) and deletion mutants (1267 bp) are indicated on each gel.
